# A genetically encoded tool to increase cellular NADH/NAD^+^ ratio in living cells

**DOI:** 10.1101/2022.09.20.508785

**Authors:** Mina L. Heacock, Evana N. Abdulaziz, Xingxiu Pan, Austin L. Zuckerman, Sara Violante, Canglin Yao, Justin R. Cross, Valentin Cracan

## Abstract

Impaired reduction/oxidation (redox) metabolism is a key contributor to the etiology of many diseases, including primary mitochondrial disorders, cancer, neurodegeneration, and aging. However, mechanistic studies of redox imbalance remain challenging due to limited strategies which can perturb cellular redox metabolism and model pathology in various cellular, tissue, or organismal backgrounds without creating additional and potentially confounding metabolic perturbations. To date, most studies involving impaired redox metabolism have focused on oxidative stress and reactive oxygen species (ROS) production; consequently, less is known about the settings where there is an overabundance of reducing equivalents, termed reductive stress. NADH reductive stress has been modeled using pharmacologic inhibition of the electron transport chain (ETC) and ethanol supplementation. Still, both these methods have significant drawbacks. Here, we introduce a soluble transhydrogenase from *E. coli* (*Ec*STH) as a novel genetically encoded tool to promote NADH overproduction in living cells. When expressed in mammalian cells, *Ec*STH, and a mitochondrially-targeted version (mito*Ec*STH), can elevate the NADH/NAD^+^ ratio in a compartment-specific manner. Using this tool, we determine the metabolic and transcriptomic signatures of NADH reductive stress in mammalian cells. We also find that cellular responses to NADH reductive stress, including blunted proliferation, are dependent on cellular background and identify the metabolic reactions that sense changes in the cellular NADH/NAD^+^ balance. Collectively, our novel genetically encoded tool represents an orthogonal strategy to perturb redox metabolism and characterize the impact on normal physiology and disease states.

## INTRODUCTION

The driving force for many critical metabolic reactions is provided by two pyridine nucleotide coenzymes: NAD(H) and NADP(H) (1–3). The redox potentials of the NADH/NAD^+^ and NADPH/NADP^+^ coenzyme couples are derived from the concentrations of their oxidized and reduced forms, which is in turn linked to concentrations of reactants and products of metabolic reactions they participate in. The cytosolic NADH/NAD^+^ pool is maintained with a large excess of oxidized NAD^+^; this makes NAD^+^ available to capture reducing equivalents from substrate oxidation reactions, primarily glycolysis and the tricarboxylic acid (TCA) cycle. Captured reducing equivalents in the form of NADH are subsequently transferred to Complex I of the electron transport chain (ETC) or used to reduce other metabolites, for example, in ketone body metabolism and fatty acid desaturation. In contrast, the cytosolic NADPH/NADP^+^ couple is maintained with a large excess of reduced NADPH. This drives reductive biosynthetic reactions such as deoxyribonucleotide and tetrahydrofolate generation, gluconeogenesis from pyruvate and the hydroxylation of fatty acids. The large excess of NADPH also supports numerous antioxidant reactions and also ROS-producing NADPH oxidases that professionally produce reactive oxygen species (ROS) in cells.

In recent years, oxidative stress has been widely studied, and we have previously reported the development of genetic tools that can be expressed in mammalian cells to induce a pro-oxidative stress through the depletion of NADH (4) or NADPH (5), respectively. Of equal or greater physiological importance are circumstances that result in an overabundance of reducing equivalents, as seen with mitochondrial dysfunction, oxygen limitation, and over-nutrition, but strategies to create reductive stress in cellular or organismal models have remained limited (6). Two methods to generate NADH reductive stress (defined as an elevated NADH/NAD^+^ ratio) are pharmacological inhibition of ETC and ethanol supplementation. Inhibiting the ETC pharmacologically blocks the ability to metabolize NADH produced by TCA cycle dehydrogenases and mitochondrial redox shuttles. This leads to an increased cellular NADH/NAD^+^ ratio, but the use of ETC inhibitors to increase mitochondrial NADH levels precludes studies of reductive stress in the context of intact oxidative phosphorylation and mitochondrial ATP generation.

Ethanol supplementation promotes reductive stress through the action of cytosolic alcohol dehydrogenases (ADHs) that convert ethanol and NAD^+^ to acetaldehyde and NADH; mitochondrial aldehyde dehydrogenases (ALDHs) can further contribute to the NADH production by further converting acetaldehyde and NAD^+^ to acetate and NADH (7–9). In a recent study, alcohol supplementation was used in mice to show that an elevated hepatic NADH/NAD^+^ ratio is linked to increased levels of 2-hydroxybutyrate (2HB), a biomarker of reductive stress associated with multiple human pathologies including mitochondrial disorders and impaired glucose tolerance (10–13). Although ethanol effectively increases the NADH/NAD^+^ ratio, overproduction of both acetaldehyde and acetate (products of ADH and ALDH catalyzed reactions) each have independent metabolic fates that are potentially confounding, and ethanol supplementation depends on expression levels of ADH and ALDH enzymes (9).

Genetic tools can provide an orthogonal strategy to perturb redox metabolism and have proven valuable because they can be expressed in an inducible fashion, targeted to subcellular compartments and expressed in a cell type-specific manner in various model systems. Previous attempts to generate reductive stress using genetic tools have been the overexpression of glucose-6-phosphate dehydrogenase (G6PD), to elevate NADPH levels, and expression of a point mutant of αB-crystallin, to elevate GSH levels (6). However, developing a genetic tool capable of overproducing NADH requires coupling to a co-substrate with more negative redox potential. For this reason, we turned our attention to bacterial **<u>s</u>**oluble **<u>t</u>**rans**<u>h</u>**ydrogenases (STHs) from the two-Dinucleotide Binding Domains Flavoprotein (tDBDF) superfamily that catalyze the interconversion of NAD^+^ + NADPH to NADH + NADP^+^ (**Fig. 1A**). STH enzymes have been previously used in multiple metabolic engineering applications in bacteria and yeast because of their ability to shuttle reducing equivalents between NADH and NADPH pools but to our knowledge have not been used to perturb redox metabolism mammalian cells (14–21). In this respect, a major advantage of STHs is that they do not require additional co-substrates and therefore do not produce or consume metabolites outside the pyridine nucleotide coenzyme couples and therefore could have less confounding effects on other metabolic pathways than other strategies that generate reductive stress (**Fig. 1A**) (22).

**Figure 1.**
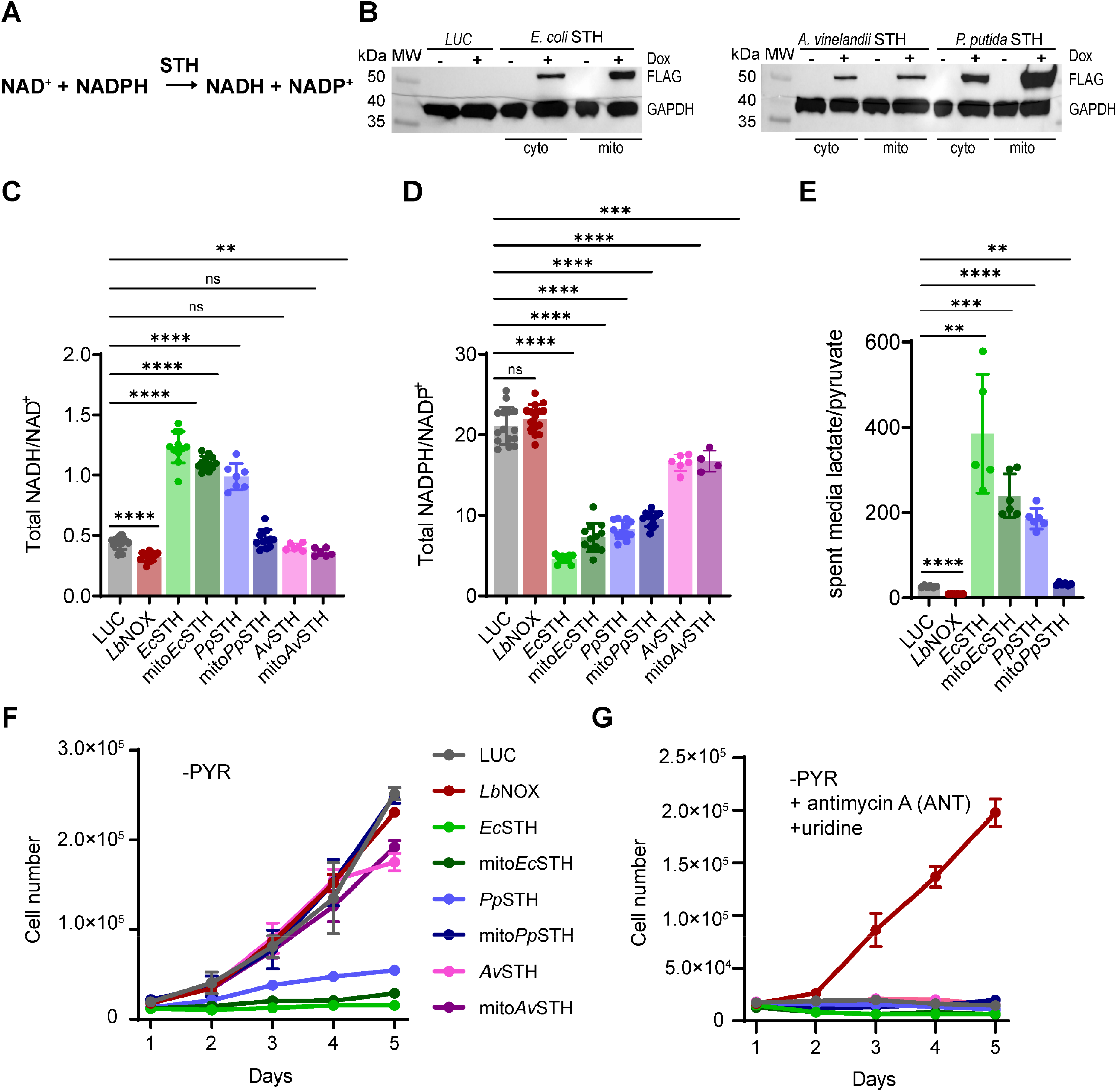
Screening of bacterial soluble transhydrogenases (STHs) for their ability to elevate NADH levels in mammalian cells. **(A)** The reaction catalyzed by a soluble transhydrogenase (STH). **(B)** Western blot analysis of HeLa cells expressing untargeted (cyto) and mitochondrially targeted (mito) bacterial STHs from *E. coli*, *A. vinelandii*, and *P. putida* under doxycycline (Dox) control. The total cellular NADH/NAD^+^ **(C)** and NADPH/NADP^+^ **(D)** ratios measured in HeLa cells expressing untargeted and mitochondrially targeted STHs from *E. coli*, *P. putida*, and *A. vinelandii*. **(E)** The lactate/pyruvate ratio measured in pyruvate-free DMEM^+dFBS^ media which was incubated for 3 hours with HeLa cells expressing untargeted and mitochondrially targeted STHs from *E. coli* and *P. putida*. **(F)** The effect of expression of untargeted and mitochondrially targeted STHs from *E. coli*, *P. putida*, and *A. vinelandii* on proliferation in pyruvate-free (-PYR) DMEM^+dFBS^. **(G)** The same as in (F) but in the presence of 1 μM antimycin A (ANT) and 200 μM uridine. LUC and *Lb*NOX expressing HeLa cells were used as controls in (B-G). The statistical significance indicated for (C-E) represents a Welch ANOVA test with unpaired t test. p<0.05*, p<0.01**, p<0.001**, p<0.0001****. For growth curves (F-G) error bars represent S.D. based on (n=3) replicates per time point per experimental condition. All cellular proliferation experiments were repeated at least n=3 times.

Here we introduce a soluble transhydrogenase from *E. coli* (*Ec*STH) as a novel genetically encoded tool to elevate the NADH/NAD^+^ ratio and induce reductive stress in mammalian cells. We show that *Ec*STH expression robustly increases the cellular NADH/NAD^+^ ratio at the expense of consuming NADPH. Using both untargeted and mitochondrially targeted *Ec*STH, we characterize metabolomic and transcriptomic signatures of the NADH reductive stress in HeLa cells. We also demonstrate that overproduction of NADH leads to the antiproliferative effect that is dependent on the cellular background. Together these data present a new genetically encoded tool and a strategy to produce a pro-reductive shift with temporal and spatial resolution. We anticipate that the new tool will be useful for investigating redox metabolism in various cellular and organismal contexts and disease states.

## RESULTS

### Bacterial soluble transhydrogenases (STHs) generate redox imbalance in mammalian cells

We initially screened three different soluble transhydrogenases (from *Escherichia coli, Azotobacter vinelandii*, and *Pseudomonas putida*) for their ability to modulate the NADH/NAD^+^ ratio when expressed in mammalian cells. Codon-optimized, C-terminal FLAG-tagged constructs, with and without a mitochondrial targeting sequence, were expressed in HeLa cells under a doxycycline (Dox)-inducible promoter (**Fig. 1B**). All experiments were performed in pyruvate-free DMEM supplemented with dialyzed FBS (DMEM^+dFBS^) to ensure exogenous pyruvate was not available to rebalance the cytosolic NADH/NAD^+^ ratio via lactate dehydrogenase (LDH)-mediated conversion of pyruvate to lactate. Under these conditions, we observed that total cellular NADH/NAD^+^ ratios were elevated by the expression of the *Ec*STH, mito*Ec*STH, and *Pp*STH transgenes (**Fig. 1C** and **Fig. S1A-C**). As controls, we included HeLa cells expressing luciferase (LUC) or a water-forming NADH oxidase from *Lactobacillus brevis* (*Lb*NOX), which we recently introduced as a genetically encoded tool to decrease cytosolic NADH/NAD^+^ ratio (4). Interestingly, the observed increase in NADH/NAD^+^ ratio with *Ec*STH expression exceeded that obtained by treating control *wild-type* HeLa cells with piercidin A, antimycin A or oligomycin A, inhibitors of mitochondrial complexes I, III or V, respectively (**Fig. S1G**). We hypothesized that the high NADPH/NADP^+^ ratio present in mammalian cells would drive bacterial STH enzymes exclusively in the direction of NADH production, at the expense of reducing equivalents from the NADPH coenzyme pool. In support of this, we observed a decrease in the whole cell NADPH/NADP^+^ ratio with expression of STH constructs (**Fig. 1D** and **Fig. S1D-F**). This observation also has physiological relevance as mitochondrial dysfunction and associated elevated NADH/NAD^+^ ratio has been linked to decreased NADPH levels (23). To further confirm that cells expressing *Ec*STH and mito*Ec*STH were experiencing reductive stress, we measured lactate and pyruvate levels in spent medium since the extracellular lactate/pyruvate ratio is a proxy for the cytosolic NADH/NAD^+^ ratio (**Fig. 1E** and **Fig. S1I-J**). This corroborated the NADH-producing activity of the three most active constructs (*Ec*STH, mito*Ec*STH, and *Pp*STH) and suggested that the LDH-mediated conversion of pyruvate to lactate is active but insufficient to rebalance the NADH/NAD^+^ ratio under these conditions.

Interestingly, the proliferation of HeLa cells expressing the three most active STH constructs was also blunted (**Fig. 1F** and **Fig. S1K**). The most parsimonious explanation for the observed anti-proliferative effect under conditions of NADH overproduction is that cells cannot maintain adequate levels of oxidized NAD^+^, a well-recognized redox requirement of proliferation (4,24,25). In support of this interpretation, the proliferation of HeLa cells expressing all STH constructs or LUC was also blunted by treatment with antimycin A (**Fig. 1G** and **Fig. S1L**). Under these conditions, only *LbNOX* expression rescued proliferation by directly catalyzing the conversion of NADH to NAD^+^ (with uridine present in the medium to bypass of coenzyme Q-dependent dihydroorotate dehydrogenase (DHODH) (**Fig. 1G** and **Fig. S1L**) (4,26). Together these results suggest that, when expressed in HeLa cells, STHs catalyze the forward NADH-producing reaction using reducing equivalents from NADPH, even when the NADH/NAD^+^ ratio is elevated by ETC inhibition. Since *Ec*STH and mito*Ec*STH generated the most pronounced increase in NADH/NAD^+^ ratio and blunted proliferation, we selected these constructs to investigate further.

### Biochemical properties of recombinant *Ec*STH

We next overexpressed the Hisx_6_-*Ec*STH-FLAG construct in *E. coli* to obtain this protein in sufficient quantities for its biochemical characterization. Purified *Ec*STH had a yellow color in solution with a characteristic UV-Visible absorption spectrum of the FAD cofactor (λ_max_ = 370 and 444 nm) (**Fig. 2A**). The molecular weight of *Ec*STH was determined to be 650 ± 30 kDa using size-exclusion chromatography and indicates that the protein is a dodecamer in solution. Enzymatic assays involving NAD(P)-dependent enzymes are typically monitored by changes in absorbance at 340 nm, but this cannot be used with STH since NADH and NADPH are spectroscopically indistinguishable. We therefore employed thio-NADH and thio-NAD^+^ analogs as thio-NADH can be monitored by absorption changes at 398 nm (ε_398_ = 11,300 M^-1^ cm^-1^) and does not overlap with 340 nm peak of NADH or NADPH. Using combinations of NADPH, NADH, NADP^+^, NAD^+^ and thio-analogs we were able to determine the kinetic parameters of *Ec*STH in both the forward and reverse directions (**Fig. 2B-C** and **Fig. S2A-D**). The determined *K_M_*’s for NAD^+^, NADPH and NADH were (68 ± 8, 32 ± 3 and 79 ± 8μM) while *k*_cat_’s were (11 ± 2, 30 ± 1 and 64 ± 7 s^-1^), respectively, indicating that neither the forward nor reverse direction are strongly favored, and the direction of the *Ec*STH catalyzed reaction will be determined by the relative abundance of the pyridine nucleotide reactants. All kinetic parameters determined for *Ec*STH in this study are summarized in (**Fig. S2D**).

**Figure 2.**
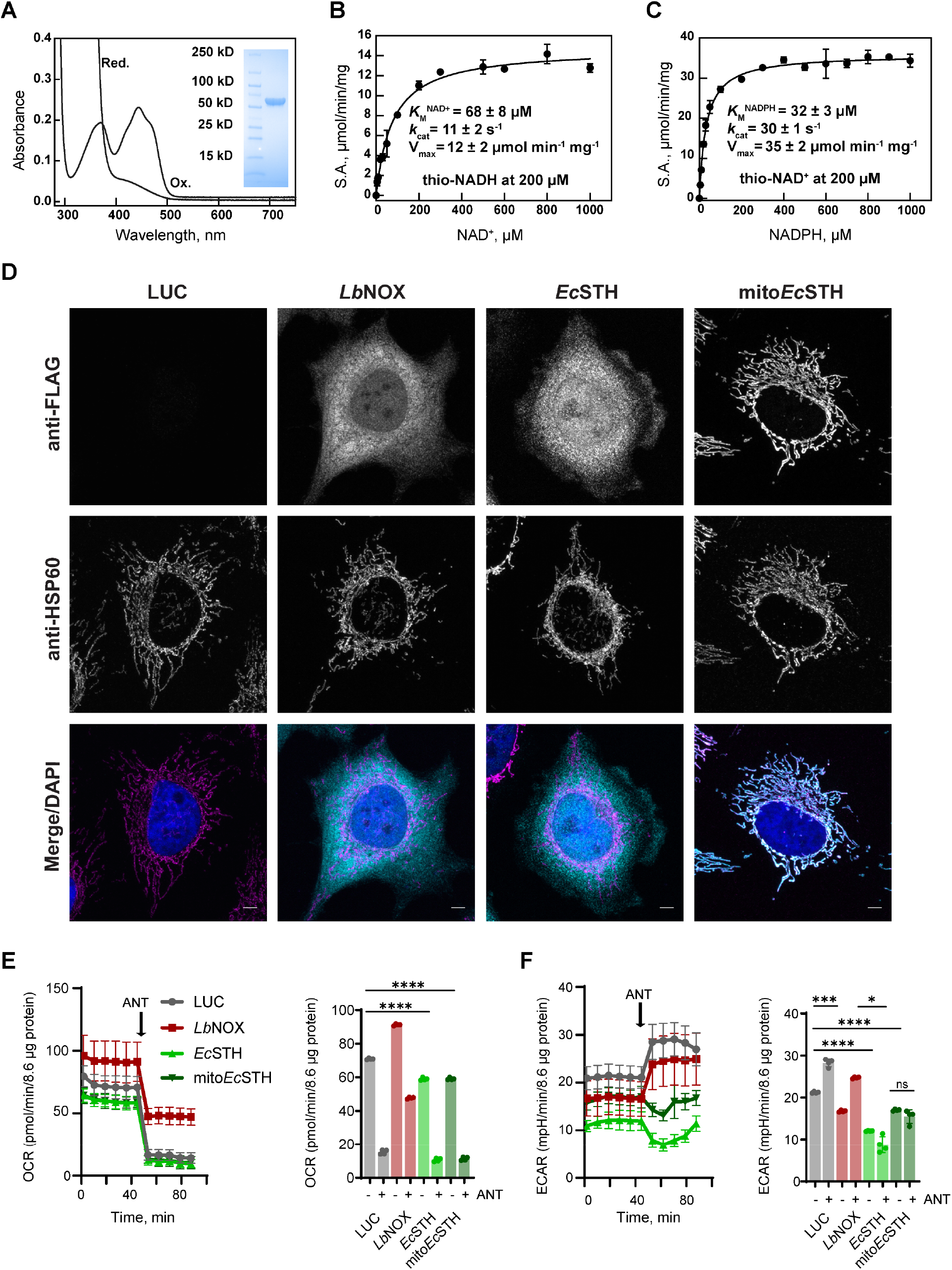
Biochemical characterization, spatial localization of *Ec*STH and mito*Ec*STH in HeLa cells and impact on oxygen metabolism. **(A)** UV-Visible spectrum of recombinant STH from *E. coli* (*Ec*STH) as purified at 20 μM FAD of active sites (Ox.) and after addition of excess of sodium dithionite (Red.). *Insert:* the SDS-PAGE of purified *Ec*STH. Michaelis-Menten analysis of the reaction catalyzed by *Ec*STH with NAD^+^ **(B)** or NADPH **(C)**. Reported values for V_max_, *k*_cat_ and *K*_M_ for NAD^+^ and NADPH represent the mean ± S.D. from (n=3) independent experiments when corresponding cosubstrates (thio-NADH or thio-NAD^+^) were fixed at 200 μM. *k*_cat_ values were calculated per monomers of FAD active sites. **(D)** Subcellular localization of untargeted *Ec*STH and mitochondrially targeted *Ec*STH (mito*Ec*STH) in HeLa cells determined by fluorescence microscopy. HeLa cells expressing FLAG-tagged *Lb*NOX, *Ec*STH, and mito*Ec*STH proteins were co-stained for FLAG tag, Hsp60 (mitochondrial marker) and DAPI (nucleus marker). Scale bars: 5 μm. Oxygen consumption rate (OCR) **(E)** and extracellular acidification rate (ECAR) **(F)** of HeLa cells expressing *Ec*STH and mito*Ec*STH before and after addition of 1 μM antimycin A (ANT) measured in pyruvate free HEPES/DMEM^+dFBS^ media. Luciferase and *Lb*NOX expressing HeLa cells were used as controls in (D-F). The statistical significance indicated for (E-F) represents a Welch ANOVA test with an unpaired t test. p<0.05*, p<0.01**, p<0.001**, p<0.0001****.

### Spatial localization of *Ec*STH and mito*Ec*STH in HeLa cells and impact on oxygen metabolism

We next used fluorescence microscopy to confirm the cellular localization of FLAG-tagged *Ec*STH and mito*Ec*STH expressed in HeLa cells (**Fig. 2D**). Untargeted *Ec*STH localized diffusely throughout the cytoplasm and the nucleus. In contrast, *LbNOX* was partially excluded from the nucleus, and mito*Ec*STH was exclusively localized in mitochondria, as evidenced by the overlap with the mitochondrial marker HSP60 (**Fig. 2D**).

Oxygen consumption rate (OCR) was mildly decreased in HeLa cells with expression of both *Ec*STH and mito*Ec*STH, indicating that increased ETC activity is not a pathway that HeLa cells can engage to relieve NADH accumulation (**Fig. 2E** and **Fig. S2E**). Mammalian cells usually respond to ETC inhibition by a compensatory increase in glycolysis, which can be measured as an increase in extracellular acidification rate (ECAR). As expected, ECAR was robustly elevated following the addition of antimycin A or oligomycin A in LUC and *Lb*NOX expressing controls, but this increase was not observed in cells expressing *Ec*STH and mito*Ec*STH (**Fig. 2F** and **Fig. S2F**). This suggests glycolysis is operating at the maximum capacity in cells expressing *Ec*STH and mito*Ec*STH, likely due to limited NAD^+^ availability, and cannot be further upregulated in response to ETC inhibition.

To investigate the impact of *Ec*STH expression on ETC-independent oxygen metabolism and the antioxidant machinery, we measured the whole cell GSH/GSSG ratios, as well as superoxide and H_2_O_2_ levels, in *Ec*STH and mito*Ec*STH expressing cells as well as in controls (**Fig. S3A-K**). In these experiments, superoxide was monitored using CellROX Green (localizes primarily to the nucleus and mitochondria) and mitoSOX (localizes primarily to the mitochondria); H_2_O_2_ levels were probed with Amplex Red (27,28). We observed that expression of *Ec*STH and mito*Ec*STH did not impact the total cellular GSH/GSSG ratio (**Fig. S3A-B**). At the same time, superoxide and H_2_O_2_ were only slightly accumulated in HeLa cells expressing *Ec*STH compared to a robust impact of menadione (vitamin K3) (**Fig. S3C-K**). These results show that, despite a sizable decrease in the total NADPH/NADP^+^ ratio, *Ec*STH and mito*Ec*STH expression do not significantly alter intracellular GSH/GSSG ratio or ROS production. This result also agrees with previous studies showing that NADPH and GSH pools are not in equilibrium (1,29).

### Metabolic features of the NADH reductive stress in HeLa cells

We next employed both untargeted and targeted metabolomics to explore how *Ec*STH and mito*Ec*STH expression impacts the metabolome of HeLa cells (**Fig. 3A-C** and **Fig. S4A-C**). In untargeted profiling, both *Ec*STH and mito*Ec*STH led to more significantly altered features than *Lb*NOX expressing cells. This suggests the cellular metabolome of HeLa cells is substantially more impacted under conditions of reductive stress compared to the pro-oxidative redox shift established by *Lb*NOX expression. One of the most significantly altered features identified was pyruvate which is illustrative of the change in the redox balance. When *Lb*NOX is expressed, the conversion of pyruvate to lactate is no longer needed to recycle NAD^+^ and hence these cells accumulate pyruvate (**Fig. 3A** and **Fig. S4A**). In contrast, cells that express *Ec*STH and mito*Ec*STH have a high NADH burden and consequently pyruvate is consumed by LDH to recover NAD^+^, resulting in dramatically reduced intracellular pyruvate levels (**Fig. 3B-C** and **Fig. S4B-C**) (24).

**Figure 3.**
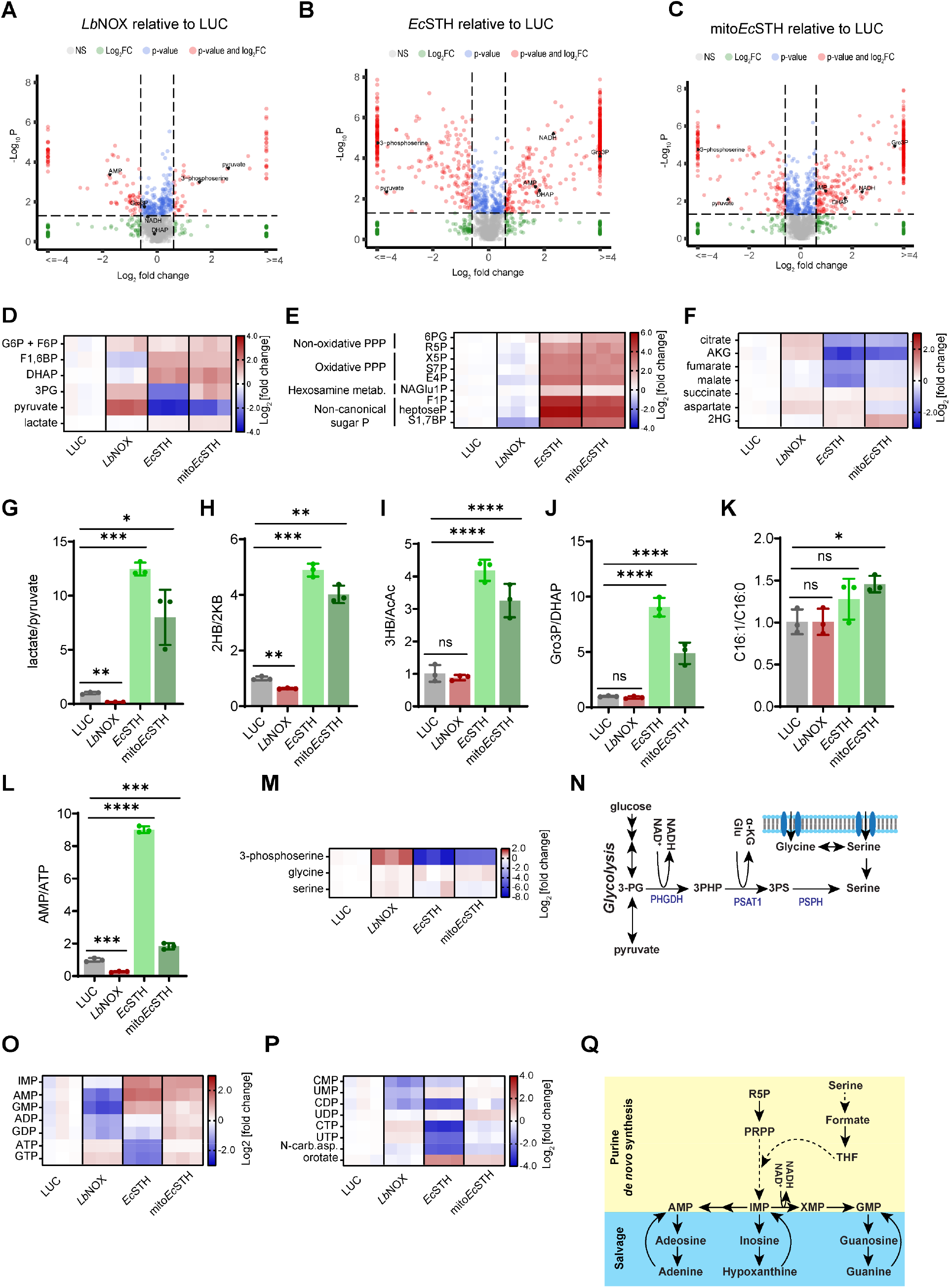
Metabolic features of the NADH reductive stress in HeLa cells. (**A-C**) Untargeted metabolomics of HeLa cells expressing *Lb*NOX, *Ec*STH and mito*Ec*STH. Heatmaps of the most impacted glycolysis (**D**), pentose phosphate pathway, non-canonical sugar phosphates (**E**) and the TCA cycle (**F**) intermediates in HeLa cells expressing *Ec*STH and mito*Ec*STH. Each column represents a biological replicate. G6P: glucose-6-phosphate; F6P: fructose-6-phosphate; F1,6BP: fructose-1,6-biphosphate; DHAP: dihydroxyacetone phosphate; 3PG: 3-phosphoglycerate; 6PG: 6-phosphogluconate; R5P: ribose-5-phosphate; X5P: xylulose-5-phosphate; S7P: sedoheptulose-7-phosphate, E4P: erythrose-4-phosphate; NAGlu1P: N-acetylglucosamine-1-phosphate; F1P: fructose-1-phosphate; heptoseP: a putative heptose with one phosphate; S1,7BP: sedoheptulose-1,7-bisphosphate; AKG: α-ketoglutarate; 2HG: 2-hydroxyglutarate. Intracellular lactate/pyruvate (**G**), 2-hydroxybutyrate/2-ketobutyrate (2HB/2KB) (**H**), 3-hydroxybutyrate/acetoacetate (3HB/AcAc) (**I**) and glycerol 3-phosphate/dihydroxyacetone phosphate (Gro3P/DHAP) (**J**), C16:1/C16:0 (**K**) and AMP/ATP (**L**) ratios measured in HeLa cells expressing *Ec*STH and mito*Ec*STH. (**M**) The heat map of intracellular levels of 3-phosphoserine, glycine and serine in *Ec*STH and mito*Ec*STH expressing HeLa cells. **(N)** Schematic of serine cellular biosynthesis. Heatmaps of the most impacted purines (**O**) and pyrimidines (**P**) in *Ec*STH and mito*Ec*STH expressing HeLa cells. N-car.asp.: N-carbamoyl-aspartate. (**Q**) Schematic of purine cellular salvage and biosynthesis. Luciferase and *Lb*NOX expressing HeLa cells were used as controls in (A-M) and (O-P). The statistical significance indicated for (A-C) represents p value cutoff = 0.05, fold change cutoff = 1.5 and Welch t test (unadjusted). The statistical significance indicated for (G-L) represents a Welch ANOVA test with an unpaired t test. p<0.05*, p<0.01**, p<0.001**, p<0.0001****.

We then turned to targeted metabolomics to specifically investigate the impact of *Ec*STH expression on NAD(H)-linked metabolic pathways. We observed that expression of *Ec*STH and mito*Ec*STH led to a robust accumulation of intermediates in upper glycolysis and non-oxidative segments of the pentose phosphate pathway (PPP) (**Fig. 3D-E**), likely due to the reduced levels of NAD^+^ limiting flux though lower glycolysis (30). In these experiments, we also noticed that *Ec*STH expression led to a dramatically increased total signal intensity of the chromatographic region where multiple sugar phosphates elute, and this increase in signal was not completely explained by metabolites typically considered to be part of upper glycolysis and the PPP (**Fig. 3E** and **Fig. S5**). Subsequently we were able to annotate some of these additional peaks as non-canonical sugar phosphates, including fructose-1-phosphate and sedoheptulose-1,7-bisphosphate, that were highly accumulated in response to *Ec*STH and mito*Ec*STH expression (**Fig. 3E**). The increase in 1-phoshate containing sugars may be explained by relaxed substrate specificity of cellular aldolases and transketolases, or by overwhelming of the DUF89 family of proteins that were recently identified as damage pre-emption enzymes acting as a disposal pathway for on non-canonical fructose phosphates formed as enzymatic side products (31,32).

Notably, *Ec*STH and mito*Ec*STH expressing cells also had lower levels of TCA cycle intermediates: citrate, aconitate, α-ketoglutarate (αKG), fumarate, and malate, compared to the LUC control (**Fig. 3F**). This result is expected as elevated NADH levels are known to downregulate TCA cycle enzymes via inhibition of pyruvate dehydrogenase complex (PDH) activity (25,33). In addition, 2-hydroxyglutarate (2HG) levels were elevated with both *Ec*STH and mito*Ec*STH, indicating that 2HG production may contribute to the cellular response to high NADH levels (**Fig. 3F**).

We then plotted the ratio of substrate/product pairs for NADH/NAD^+^-linked reactions. Many of these are equilibrium reactions and can therefore provide a surrogate for the NAD(H) pool redox status and identify metabolic reactions used by cells to counter reductive stress driven by NADH overaccumulation (**Fig. 3G-K**). In agreement with the changes in spent culture media, the intracellular lactate/pyruvate ratio was substantially elevated in *Ec*STH and mito*Ec*STH expressing cells (**Fig. 3G** and **Fig. S6A-B**) (34,35). 2-hydroxybutyrate (2HB, also known as alpha-hydroxybutyrate) is produced from 2-ketobutyrate (2KB) downstream of threonine, homocysteine, and methionine metabolism and was recently reported to be a biomarker of NADH reductive stress in mitochondrial encephalomyopathy lactic acidosis and stroke-like episodes (MELAS), and other conditions linked to redox imbalance (34,35). Two enzymes, LDH and α–hydroxybutyrate dehydrogenase, catalyze the interconversion of 2KB and 2HB, and both are NADH/NAD^+^-linked reactions (11). As expected, we observed a substantial increase in 2HB and a decrease in 2KB levels in HeLa cells expressing *Ec*STH and mito*Ec*STH and this resulted in a substantially elevated 2HB/2KB ratio (**Fig. 3H** and **Fig. S6C-D**). Similar changes in 2HB and 2KB were observed in *wild-type* HeLa cells treated with different ETC inhibitors, with the most pronounced impact in the presence of antimycin A (**Fig. S6R-T**). Another redox pair that participates in the buffering of the cellular redox environment is an equilibrium reaction between 3-hydroxybutuyrate (3HB) and acetoacetate (AcAc), catalyzed by 3-hydroxybutyrate dehydrogenase and considered a proxy for the mitochondrial NADH/NAD^+^ ratio (36). We observed that *Ec*STH expression did not substantially impact 3HB levels but AcAc levels were drastically reduced by expression of both *Ec*STH and mito*Ec*STH, resulting in a significantly increased 3HB/AcAc ratio (**Fig. 3I** and **Fig. S6E-F**).

In principle, the glycerol-3-phosphate (Gro3P) shuttle system is also an important vent for NADH surplus. This enzyme system converts the glycolytic intermediate dihydroxyacetone phosphate (DHAP) to Gro3P, and by doing so regenerates NAD^+^ from NADH (30). Subsequently, Gro3P is converted back to DHAP and electrons are donated to the mitochondrial coenzyme Q pool. Evidence that the Gro3P shuttle is intact in *wild-type* HeLa cells was obtained by treating cells with antimycin A to block progression of electrons beyond the coenzyme Q pool to cytochrome c. This led to a 65-fold accumulation of Gro3P (**Fig. S6M-O)**. In response to *Ec*STH expression, and to a lesser extent with mito*Ec*STH, we also observed a robust accumulation of both Gro3P and DHAP and a substantially elevated Gro3P/DHAP ratio (**Fig. 3J** and **Fig. S6G-H**). However, it is important to note that in HeLa cells the Gro3P shuttle was not able to successfully vent excess of NADH through the coenzyme Q pool and ETC, as evidenced by the decreased oxygen consumption and reduced proliferation. Another NADH-consuming mechanism that was recently reported to relieve NADH-induced reductive stress is fatty acid desaturation (37) and in cells expressing mito*Ec*STH we observed an increase in C16:1/C16:0 ratio (**Fig. 3K** and **Fig. S6I, J**). Aspartate levels have been directly impacted by the ETC activity and associated changes in cellular NADH/NAD^+^ (38). We observed that a blockade of ETC in *wild-type* HeLa cells diminished aspartate levels, but *Ec*STH and mito*Ec*STH expression only slightly impacted aspartate levels compared to LUC or *Lb*NOX controls (**Fig. S6P-Q)**. These later results also highlight the importance of separating reductive stress from mitochondrial dysfunction.

One of the most dramatically diminished metabolites in response to *Ec*STH and mito*Ec*STH expression was 3-phosphoserine, while glycine and serine levels were only modestly impacted (**Fig. 3M-N**). This can be explained by 3-phosphoglycerate dehydrogenase (PHGDH), a key enzyme in the serine biosynthesis pathway, having a *K*_d_ for NADH of 0.22 ± 0.03 μM, which is >2000-fold lower compared to a *K*_d_ for NAD^+^ (444 ± 18 μM); therefore, under conditions of NADH reductive stress, serine biosynthesis from PHGDH cannot proceed in the forward direction (39–41). Intracellular serine and glycine levels were however not significantly impacted, likely because intracellular serine and glycine can both supported by serine in the culture media (**Fig. 3N**). In addition to the already highlighted changes in central carbon metabolism, we also observed a significant impact on the levels of purines and pyrimidines (**Fig. 3L, O-P** and **Fig. S6K-L**). Since HeLa cells expressing *Ec*STH and mito*Ec*STH are not proliferating, it is not surprising that AMP accumulated, while ATP levels diminished. Accumulation of purines such as IMP, AMP and GMP may be explained by elevated levels of ribose 5-phosphate in these cells, as the conversion of ribose 5-phosphate to phosphoribosyl pyrophosphate (PRPP) is the first committed step in IMP and purine biosynthesis (**Fig. 3O, Q**)(41).

Collectively our data suggest that the NADH reductive stress in HeLa cells leads to extensive metabolic remodeling that includes upregulation of non-oxidative PPP, activation of purine biosynthesis, and a blockade of serine biosynthesis from 3-phosphoglycerate.

### Transcriptomic signatures of the NADH reductive stress in HeLa cells

To further explore how the NADH reductive stress impacts cellular metabolism, we performed RNA sequencing (RNA-Seq) analysis of HeLa cells expressing luciferase, *Lb*NOX, *E*cSTH or mito*Ec*STH for 24 hours. Gene expression in cells expressing *Lb*NOX changed modestly when compared with the luciferase control (**Fig. 4A**). By contrast, there are more than 600 upregulated and more than 700 downregulated genes identified in cells expressing *EcSTH* (**Fig. 4B**). This mirrors the metabolomic conclusions and indicates that cells undergo a larger transcriptomic response to NADH reductive stress than to a pro-oxidative shift in redox metabolism. Similar results were obtained in cells expressing mito*Ec*STH (**Fig. 4C**). Among the most significantly changed genes were the growth differentiation factor 15 (*GDF15*) (upregulated), while the pregnancy zone protein (*PZP*) was downregulated in both *EcSTH* and mito*Ec*STH expressed cells. Consistently, *GDF15* was recently suggested as a biomarker of the NADH reductive stress linked to mitochondrial disease (35), while *PZP* is a newly characterized biomarker for lung adenocarcinoma (LAC) or colorectal cancer (CRC) in type 2 diabetes mellitus patients (42,43). A substantial number of genes were also differentially changed in mito*Ec*STH expressing cells compared with *EcSTH* (**Fig. 4D**), suggesting the cellular response to the NADH reductive stress is different depending on cellular compartment. Gene ontology (GO) enrichment analysis revealed the most significantly changed processes to be “oxidation-reduction process” and “cholesterol biosynthesis process” (**Fig. 4E-G**). Interestingly, most genes in the “cholesterol biosynthesis” cluster are transcriptionally downregulated in *EcSTH*-expressing cells, such as *HMGCR, MVK, MVD, IDI1, FDPS, FDFT1*, and *SQLE*, all of which encode enzymes of the cholesterol biosynthesis pathway. The only gene upregulated in this cluster is *NPC1L1*, a cholesterol transporter in the plasma membrane that mediates cholesterol efflux into cells (44). Moreover, *ABCA13*, another gene linked to cholesterol trafficking, was one of the most upregulated genes in the *EcSTH* expressing cells (45). In a recent study where the ketogenic 3HB diet was used to induce reductive stress in mice bearing PDAC allograft tumors, the cholesterol biosynthesis was also downregulated as determined by RNA-Seq analysis, with 3 genes in common between these studies: *SQLE, FDPS* and *MSMO1* (46).

**Figure 4.**
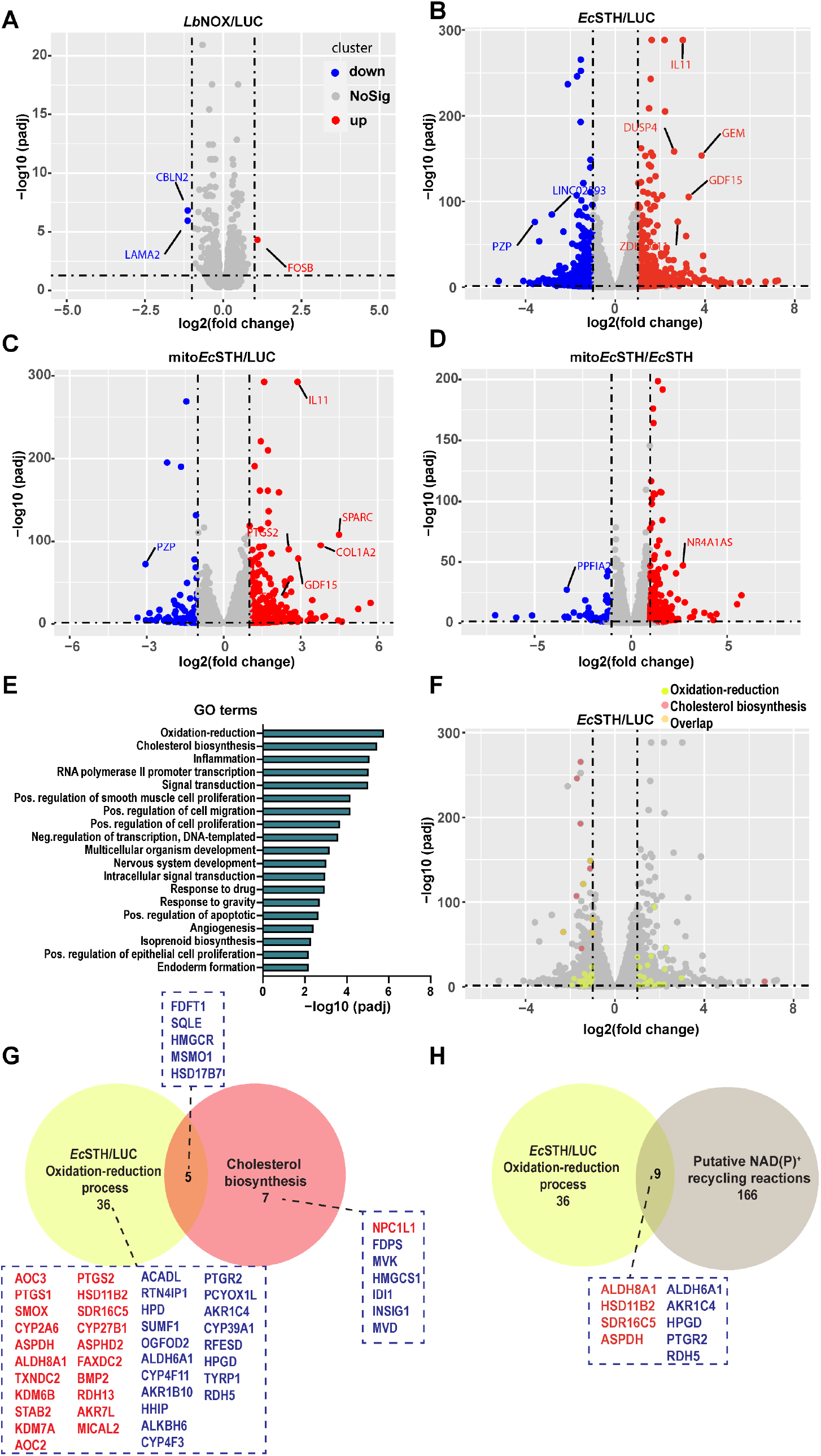
RNA-Seq of HeLa cells expressing *Lb*NOX, *Ec*STH and mito*Ec*STH. **(A-D)** Volcano plots that represent the indicated group’s log2 fold change (x-axis) and adjusted P-value for significance (y-axis). Red dots represent up-regulated genes. Blue dots represent down-regulated genes. Gray dots represent genes without significant changes in expression. Selected genes are marked with gene names. **(E)** Gene ontology enrichment analysis of *Ec*STH vs luciferase control group. **(F)** Volcano plot representing the top 2 GO terms in (E). Light green dots represent the “oxidation-reduction process” and salmon dots represent the “cholesterol biosynthetic process”. Overlapping genes are highlighted in orange. **(G-H)** Venn diagram displaying the similarities and differences of genes from the indicated gene sets. Yellow circle includes all genes in the GO term “oxidation-reduction process”. Salmon circle contains all genes in the GO term “cholesterol biosynthetic process”. Light brown circle indicates the NAD(P)ome (genes in humans that encode NAD^+^ or NADP^+^ dependent enzymes as defined in (30)). The up-regulated genes are marked in red. The down-regulated genes are marked in blue.

There are also still multiple human genes that do not have an established cellular function or known co-substrate, despite being classified as NAD(P)(H) consuming or producing (34). We find that from the “oxidation-reduction process” gene set, nine genes encode enzymes that have clear dependence on NAD(H) or NADP(H) as co-substrates; these genes were recently nominated as possible NAD(P)^+^ recycling reactions (**Fig. 4H**)(30). ALDHs and other enzymes in this group are poorly biochemically studied dehydrogenases that use retinoic acid, gamma-aminobutyric acid, or betaine as a cosubstrate (47,48). Interestingly, ALDH8A1 expression was shown to be induced by ethanol in the liver, and this enzyme catalyzes the reaction of 2-aminomuconate semialdehyde and NAD^+^ to produce 2-aminomuconate and NADH in the kynurenine pathway of tryptophan catabolism (49,50). In summary, our RNA-Seq experiments identified genes that are impacted by the NADH reductive stress induced by the novel genetically encoded tool, *Ec*STH. Our results are consistent with a recent study that showed the downregulation of cholesterol biosynthesis pathway as one of the key transcriptomic features of the NADH reductive stress (46). In addition, we identified previously uncharacterized NAD(P)-linked reactions that are important for cellular redox homeostasis and respond to an elevated cellular NADH/NAD^+^ reduction potential (**Fig. 4H**).

### Direct NAD^+^ recycling is sufficient to rescue the anti-proliferative effect of *Ec*STH expression

We next asked if the anti-proliferative effect we observed in HeLa cells expressing *Ec*STH and mito*Ec*STH is specific to the cellular background. We expressed *Ec*STH and mito*Ec*STH in C2C12 mouse myoblasts, human primary fibroblasts IMR-90 and A549, HEK-293T, MIA PaCa-2 and PANC-1 cancer cell lines under Dox control (**Fig. S7A, E, I, M, Q and U)**. Expression of *Ec*STH and mito*Ec*STH elevated the NADH/NAD^+^ and decreased the NADPH/NADP^+^ ratios to different extents in each of these cell lines, with the largest effect observed in IMR-90 fibroblasts (**Fig. S7B-C, F-G, J-K, N-O, R-S** and **V-W**). Interestingly, the impact of *Ec*STH and mito*Ec*STH expression on proliferation was most pronounced in IMR-90 fibroblasts (**Fig. S7D, H, L, P, T** and **X**). Our experiments suggest that cells vary in their ability to withstand the NADH reductive stress and that the observed total NADH/NAD^+^ ratio can be used as a predictor of the severity of the anti-proliferative effect.

We next measured whole-cell NADH/NAD^+^ and NADPH/NADP^+^ ratios in HeLa cells expressing *Ec*STH and mito*Ec*STH treated with piericidin A, antimycin A or pyruvate (**Fig. 5A-B**). We observed that both STH constructs increased the NADH/NAD^+^ ratio to a greater extent when used in combination with ETC inhibitors, with the largest effect observed for *Ec*STH combined with antimycin A treatment (**Fig. 5A**). In contrast, the NADPH/NADP^+^ ratio was not substantially impacted across all conditions (**Fig. 5A-B**) (**Fig. S1H**) and treatment with 1 mM pyruvate alleviated the build-up of NADH, while having no effect on NADPH/NADP^+^ (**Fig. 5A-B**). This suggests the *Ec*STH and mito*Ec*STH constructs can also be employed in conjunction with ETC inhibition to further amplify the NADH/NAD^+^ ratio increase.

**Figure 5.**
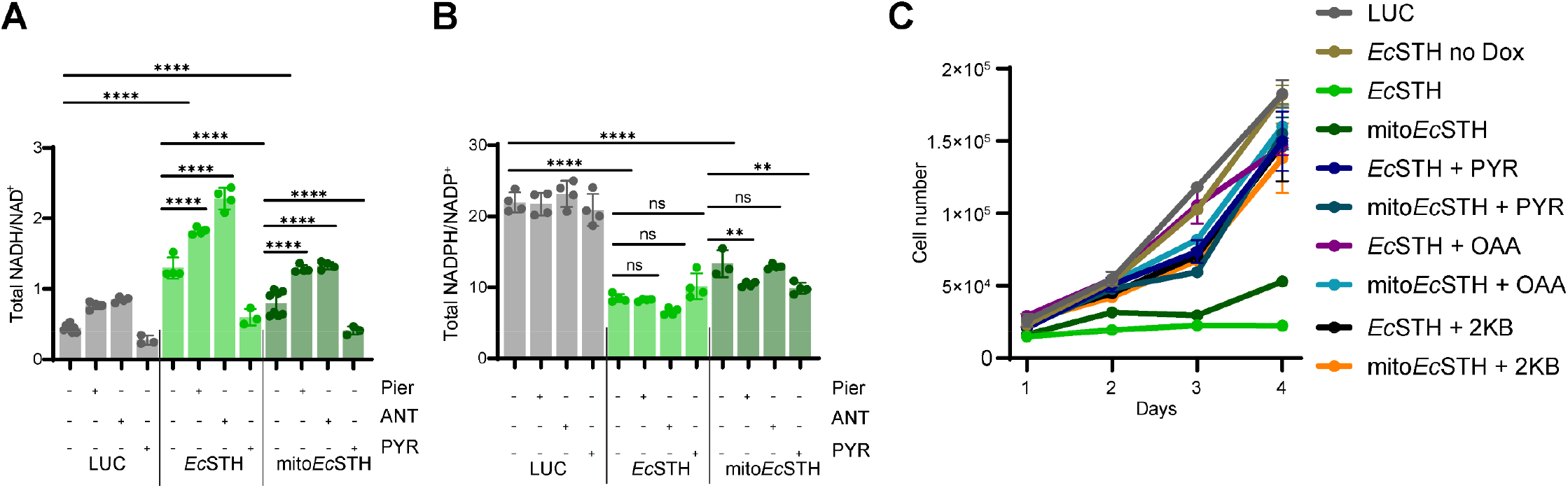
Direct NAD^+^ recycling is sufficient to rescue the anti-proliferative effect in HeLa cells. The total NADH/NAD^+^ **(A)** and NADPH/NADP^+^ **(B)** ratios measured in HeLa cells expressing *Ec*STH and mito*Ec*STH three hours after changing to fresh pyruvate-free DMEM^+dFBS^ with 1 μM antimycin A (ANT), 1 μM piericidin A (Pier), or 1 mM pyruvate (PYR). (**C**) Rescue of the anti-proliferative effect of *Ec*STH and mito*Ec*STH expression in HeLa cells when DMEM^+dFBS^ was supplemented with exogenous electron acceptors (10 mM pyruvate (PYR), 10 mM oxaloacetate (OAA) or 10 mM α-ketobutyrate (2KB)). The statistical significance indicated for (A-B) represents a Welch ANOVA test with an unpaired t test. p<0.05*, p<0.01**, p<0.001**, p<0.0001****. For growth curves (C) error bars represent S.D. based on n=3 replicates per time point per experimental condition. All cellular proliferation experiments were repeated at least n=3 times.

Finally, we tested if the proliferation defect observed in HeLa cells was primarily due to inadequate NAD^+^ recycling by exposing cells to several exogenous electron acceptors known to support NAD^+^ recycling in mammalian cells (4,38). In these experiments we observed that supplementation of the growth medium with pyruvate, 2KB, or oxaloacetate (OAA) completely rescued the anti-proliferative defect (**Fig. 5C**). This suggests that, in cells with an intact ETC, providing an alternative mechanism for NAD^+^ recycling is sufficient to rescue the anti-proliferative effect caused by *Ec*STH or mito*Ec*STH expression.

## DISCUSSION

Recent advancements in high-resolution mass spectrometry have increased our understanding of metabolism in normal physiology and disease. Hundreds of cellular metabolites can now be measured in parallel with unprecedented sensitivity (51,52), but comparatively few techniques are available to perturb central metabolic parameters with compartment-specific precision (2,53). This study combines synthetic biology and metabolomics approaches to develop and validate a novel genetically encoded tool as a new strategy to model NADH reductive stress in mammalian cells. Our approach uses bacterial soluble transhydrogenases (STHs), whose physiological function is to maintain energy metabolism and redox cofactors homeostasis (15,17,19,20,54). Unlike genetically encoded sensors that allow visualization of changes in metabolite concentrations or biochemical events to be detected based on changes in fluorescence or luminescence, genetically encoded tools are enzymes that directly impact metabolite steady-state concentrations (53). We previously used a water-forming NADH oxidase from *L. brevis* (*Lb*NOX), as well as a variant that we engineered to be strictly NADPH specific (TPNOX), to decrease cellular NADH/NAD^+^ or NADPH/NADP^+^ ratios, respectively (4,5). Here, we present the development and validation of *E. coli* soluble transhydrogenase (*Ec*STH), as a third genetically encoded tool to model the NADH reductive stress in living cells by increasing the cellular NADH/NAD^+^ ratio (**Fig. 1A**). We believe that *Ec*STH, together with *Lb*NOX and TPNOX, this work will expand the toolkit of reagents available to the scientific community to study the crosstalk between cellular NADH and NADPH pools (4,5,53).

The most widely used assay to study the reaction catalyzed by recombinant STHs employs thio-analogs of redox cofactors (54,55). This allows to monitor the STH catalyzed reaction at 398 nm (ε_398_= 11.9 mM^-1^ cm^-1^) as both NADH and NADPH cofactors absorb at 340 nm. However, it must be stressed that while thio-NADP(H) are suitable analogs of NADP(H) cofactors, thio-NAD(H) are poor NAD(H) mimetics (56). In general, thio-analogs are used as donors or acceptors of electrons irrespective of the cofactor identity, as for example thio-NADH and NAD^+^ form a productive substrate pair (**Fig. S2D**). Very similar *k*_cat_/*K*_M_ values for all reactions studied, and no activity with NADP^+^ and thio-NADH pair of substrates are in agreement with previous reports that *Ec*STH catalyzes the NADH-producing reaction when NADPH levels are high (**Fig. 2D**)(57). Indeed, in our experiments in mammalian cells *Ec*STH catalyzes the reaction exclusively in the direction of transferring reducing equivalents from NADPH to NAD^+^ (in cells with both intact or inhibited ETC) (**Fig. 1A**, **Fig. 1C** and **Fig. 5C**). This directionality is driven by the cellular state of NADPH/NADP^+^ reduction potential, as the majority of cellular NADPH is in the reduced form (36). With TPNOX and mitoTPNOX expression in HeLa cells we have previously demonstrated that mitochondrial pyridine nucleotide pools are asymmetrically connected, as oxidation of mitochondrial NADPH leads to oxidation of mitochondrial NADH, but not *vice versa*, as only mitoTPNOX was capable of rescuing pyruvate auxotrophy (5). Here we show that neither *Ec*STH nor mito*Ec*STH rescued pyruvate auxotrophy. This suggests that although mito*Ec*STH consumes NADPH, it simultaneously generates NADH making regeneration of NAD^+^ via crosstalk between mitochondrial pyridine nucleotide pools infeasible (**Fig. 1G** and **Fig.S1L**). This is supported by the additive effect in the NADH/NAD^+^ ratio observed in the presence of ETC inhibitors, particularly antimycin A (**Fig. 5A**).

Our data indicate that *Ec*STH promotes a robust increase in the NADH/NAD^+^ ratio in mammalian cells, with a clear impact on lactate/pyruvate, 2HB/2KB, and 3HB/AcAc redox pairs (**Fig. 3G-I)**. The NADH reductive stress impedes lower glycolysis, TCA cycle, and serine biosynthesis and forces carbon flux into the non-oxidative PPP as evidenced in our metabolomic profiling in HeLa cells (**Fig. 3A-F, M** and **Fig. S4A-C**). The effect on the non-oxidative PPP is also observed with TPNOX expression (5), but the impact is much less pronounced than seen with *Ec*STH expression. We also observed the upregulation of purine biosynthesis as one of the main features of reductive stress (**Fig. 3O, Q**). On the other hand, the extent to which different cell lines show a proliferation defect in response to *Ec*STH expression was linked to their ability to recycle NAD^+^ (**Fig. S7A-X**). We also find that while a pro-oxidative shift (a decrease in NADH/NAD^+^ ratio) is essentially transcriptionally silent, there is a drastic transcriptional response to the elevated NADH/NAD^+^ ratio in HeLa cells (**Fig. 4A-D**). Both our metabolomics and transcriptomics data is in alignment with prior predictions that an excess of reducing equivalents can be more deleterious for cells compared to a pro-oxidative shift (58).

Interestingly, one of the features of the transcriptional response of *Ec*STH expression was the downregulation of the cholesterol biosynthesis pathway, which was also recently identified in PDAC tumors in mice that received a keto diet (oxidation of 3HB led to the NADH reductive stress in these animals) (46). Another prominent feature of the NADH reductive stress was the upregulation of ALDH enzymes, some of which were already known as whose expression is activated as a response to ethanol-invoked reductive stress (50). This new genetic tool also allows the generation of reductive stress in a compartmentalized manner, and under conditions where ETC activity remains intact. We also hypothesize that modulation of purine biosynthesis can be one direction to counteract redox imbalance. Another interesting implication of our work is that reductive stress can also result in the accumulation of non-canonical sugar phosphates, which are known to be toxic (59).

Collectively our data describes the development and validation of *Ec*STH as a genetically encoded tool to model NADH reductive stress in living cells and report both transcriptomic and metabolomic signatures of the reductive stress. We show that cells with an intact respiratory chain vary in their ability to withstand reductive stress, possibly by providing alternative compensatory biological pathways for NAD^+^ recycling. In addition, we show that under the NADH reductive stress, non-oxidative PPP and purine biosynthesis are upregulated. Our study sets the stage for developing a technology that will allow researchers to directly test how various cells or tissues respond to reductive stress.

## MATERIALS AND METHODS

### Reagents and Supplies

All reagents and supplies were purchased from ThermoFisher Scientific, Sigma-Aldrich, and VWR unless otherwise specified.

### Cell Lines and Cell Culture

HeLa, HEK293T, and C2C12 cells were purchased from the American Type Culture Collection (ATCC), human fibroblasts IMR-90 were a gift from the Adams Laboratory (SBP, La Jolla, CA), A549 cells were a gift from the Lamia Laboratory (TSRI, La Jolla, CA), MIA PaCa-2 and PANC-1 cells were a gift from the Commisso Laboratory (SBP, La Jolla, CA). All cell lines were cultured in DMEM^+FBS^ [Dulbecco’s Modified Eagle Medium without pyruvate and glucose (ThermoFisher, 11966025) supplemented with 25 mM glucose (ThermoFisher, A2494001), 10% non-dialyzed FBS (Sigma, F2442) and 1% penicillin/streptomycin (ThermoFisher, 15140148)] at 37°C in an atmosphere of 5% CO_2_ (IMR-90 human fibroblasts were cultured in an atmosphere of 5% CO_2_/3.5% O_2_). The Tet-On 3G system lentiviral-infected HeLa and C2C12 cell lines were cultured in DMEM^+FBS^ supplemented with 0.5 mg/mL geneticin (ThermoFisher, 10131035) and 10 μg/mL puromycin (ThermoFisher, A1113803). The Tet-One-Puro system lentiviral-infected IMR-90, A549, HEK293T, MIA PaCa-2, and PANC-1 cells were cultured in DMEM^+FBS^ in the presence of 1 μg/mL puromycin. All experiments were performed in DMEM^+dFBS^ without antibiotics [pyruvate free DMEM (ThermoFisher, 11966025) supplemented with 25 mM glucose and 10% dialyzed FBS (ThermoFisher, 26400044] and other media components or chemical compounds as specified. All cell lines in this study were mycoplasma free.

### Cloning of Soluble Bacterial Transhydrogenases into the pLVX-TRE3G Vector

*Homo sapiens* codon-optimized genes encoding soluble transhydrogenases (STHs) from *Escherichia coli* (HAP3137380.1), *Pseudomonas putida* (WP_012313573), and *Azotobacter vinelandii* (WP_012700102) flanked by NotI and MluI restriction sites were custom synthesized by GENEWIZ. All constructs were delivered in the pUC57 vector and included, in addition to the original ORF: *i*) an N-terminal mitochondrial targeting sequence (MTS) of subunit IV of human cytochrome c oxidase and (*ii*) a C-terminal linker sequence with a FLAG-tag. After digestion with NotI and MluI, inserts were directly ligated into the pLVX-TRE3G vector (Clontech, CA). To remove the MTS and produce constructs of untargeted STHs, corresponding constructs were amplified using the following primers containing the NotI and MluI restriction sites which are underlined: *Escherichia coli* 5’-TTA ATT <u>GCG GCC GC</u> ATG CCC CAC AGC TAC GAT TAC G −3’; *Pseudomonas putida* 5’-TTA ATT <u>GCG GCC GC</u> ATG GCC GTG TAC AAC TAT GAC GTG −3’; *Azotobacter vinelandii* 5’-TTA ATT <u>GCG GCC GC</u> ATG GCC GTG TAC AAT TAT GAC GTG G −3’ and the reverse primer 5’-TTA ATT <u>ACG CGT TTA</u> CTT GTC ATC GTC ATC CTT GT −3’. After digestion with NotI and MluI, the PCR products were ligated into the pLVX-TRE3G vector (Clontech, CA) and verified by DNA sequencing (Eton Bioscience, San Diego, CA).

### Cloning of Luciferase, *LbNOX, Ec*STH, mito*Ec*STH into pLVX-TetOne-Puro Vector

To re-clone *Ec*STH, mito*Ec*STH and *Lb*NOX into the pLVX-TetOne-Puro vector, corresponding constructs were amplified from corresponding pLVX-TRE3G vectors using the following primers containing the EcoRI and AgeI restriction sites, which are underlined: *Ec*STH 5’-TTA ATT <u>GAA TTC</u> ATG CCC CAC AGC TAC GAT TAC G −3’; mito*Ec*STH 5’-TTA ATT <u>GAA TTC</u> ATG CTG GCC ACC AGG GTC TTT AGC C −3’; *Lb*NOX 5’-TTA ATT <u>GAA TTC</u> ATG AAG GTC ACC GTG GTC GGA TGC −3’ and the reverse primer 5’-TTA ATT <u>ACC GGT</u> TTA CTT GTC ATC GTC ATC CTT GTA ATC C −3’. To re-clone Luciferase into the pLVX-TetOne-Puro vector following primers we used containing the EcoRI and AgeI restriction sites, which are underlined: 5’-TTA ATT <u>GAA TTC</u> ATG GAA GAC GCC AAA AAC ATA AAG −3’ and 5’-TTA ATT <u>ACC GGT</u> TTA CAA TTT GGA CTT TCC GCC CTT C −3’. After digestion with EcoRI and AgeI the PCR products were ligated into the pLVX-TetOne-Puro vector (Addgene plasmid #124797).

### Cloning of *Ec*STH into pET30a Vector for Bacterial Expression

*E. coli sth* gene was amplified from the pLVX-TRE3G vector (*H. sapiens* codon-optimized sequence) using the following primers containing the BamHI and XhoI restriction sites which are underlined: 5’-TTA ATT <u>GGA TCC</u> ATG AAG GTC ACC GTG GTC GG-3’ and 5’-TTA ATT <u>CTC GAG</u> TCA CTT GTC ATC GTC ATC C-3’. After digestion the PCR product was ligated into the pET30a vector (EMD Millipore). The resulting construct encodes *Ec*STH with both an N-terminal Hisx6-tag and a C-terminal FLAG-tag.

### Lentiviral Production and Infection

Lentiviruses were produced by transfecting packaging vectors psPAX2 (psPAX2 was a gift from Didier Trono, Addgene plasmid # 12260) and pMD2.G (pMD2.G was a gift from Didier Trono, Addgene plasmid # 12259) together with (*i*) pLVX-Tet3G regulator vector (encodes the Tet-On element), (*ii*) response vector pLVX-TRE3G-Luciferase, *Lb*NOX, *Pp*STH, mito*Ep*STH, *Ec*STH, mito*Ec*STH, *Av*STH or mito*Av*STH, or (*iii*) response vector pLVX-Tet-One-Puro-Luciferase, *Lb*NOX, *Ec*STH and mito*Ec*STH into HEK293T cells as previously described (4,5). Forty-eight hours after transfection media containing the virus was collected, centrifuged, and stored at −80 °C. HeLa and C2C12 Tet3G cells were produced first using a lentivirus from the pLVX-Tet3G vector (encodes the Tet-On element), and subsequently following selection with 0.5 mg/mL of geneticin, cells were infected with lentivirus produced using pLVX-TRE3G-Luciferase, *Lb*NOX, *Ec*STH, mito*Ec*STH, *Pp*STH, mito*Ep*STH, *Av*STH, or mito*Av*STH vectors as previously described (5). IMR-90, A549, HEK293T, MIA PaCa-2, and PANC-1 TetOne cells were produced by a single infection with corresponding pLVX-TetOne-Puro-Luciferase, *Lb*NOX, *Ec*STH or mito*Ec*STH lentiviruses. The pLVX-TetOne-Puro system encodes both the Tet-On element and the response gene within a single plasmid. All cell lines were maintained in DMEM^+FBS^ in the presence of 0.5 mg/mL geneticin and 10 μg/mL puromycin (HeLa and C2C12 Tet-On 3G) or in DMEM^+FBS^ in the presence of 1 μg/mL puromycin (IMR-90, A549, HEK293T, MIA PaCa-2 and PANC-1 Tet-One).

### Immunoblots

Two hundred thousand cells were seeded in a 6-well plate in 2 mL of DMEM^+FBS^. Twenty-four hours later media was exchanged with 3 mL of DMEM^+dFBS^ ± 300 ng/mL doxycycline (Dox). After additional twenty-four hours, cells were washed with ice-cold PBS and were lysed with 300 μL of 50 mM HEPES, pH 7.5, 100 mM NaCl, 2 mM EDTA and 1% Triton X-100. Protein concentration was determined using the Bradford Assay (Bio-Rad, #5000205). After SDS-PAGE proteins were transferred using a Bio-Rad Trans-Blot Turbo transfer system. Anti-FLAG (Cell Signaling, #2368S; 1:500-1000), and anti-GAPDH (Cell Signaling, #2118L; 1:10000) were used as primary antibodies. Anti-Rabbit (Cytiva NA9340; 1:2000) was used as a secondary antibody. Membranes were imaged using a Bio-Rad ChemiDoc MP gel-documentation system.

### Immunofluorescence and Imaging

Ten thousand cells were seeded on a coverslip in a 6-well plate in 2 mL of DMEM^+FBS^. Twenty-four hours later media was exchanged to 2 mL of DMEM^+dFBS^ ± 300 ng/mL doxycycline (Dox). After an additional twenty-four hours, cells were fixed with 4% PFA (paraformaldehyde) for 15 minutes and washed 3 times with PBS for 5 minutes each. Cells were then permeabilized with 0.2% Triton X-100 in PBS for 5 minutes, followed by 3 washes in PBST (0.05% Tween20 in PBS) for 5 minutes each, and incubated with blocking buffer (0.2% BSA in PBST) for 20 minutes at room temperature. Cells were incubated with primary antibodies in blocking buffer overnight at 4 °C. The next day, cells were washed with PBST 3 times for 5 minutes each and incubated with secondary antibodies in blocking buffer for 2 hours at room temperature. After 2 hours, cells were again washed 3 times with PBST in 5 minutes increments and rinsed with 2 mL of Milli-Q water, airdried and mounted. The following primary antibodies were used: anti-FLAG (Cell Signaling, #2368S; 1:400), anti-HSP60 (Cell Signaling, #12165; 1:2000). Alexa Fluor 488- and Alexa Fluor 568-conjugated secondary antibodies were used (ThermoFisher, A-11034, A-11036). VECTASHIELD mounting medium with DAPI (Vectorlabs, H-1000) was used to visualize nuclei. Cells were imaged using a Leica TCS SP8 confocal laser scanning microscope with HC Plan APO 63x NA 1.4 oil objective. All images were analyzed in Fiji (60).

### Proliferation Assays

Five thousand or ten thousand cells were seeded in 0.5 mL of DMEM^+FBS^ in 24-well plates. The next day media was exchanged with 1 mL of DMEM^+dFBS^ supplemented with 300 ng/mL of Dox and other components as indicated. Proliferation assays with IMR-90 cells were always performed at 3.5% oxygen. On days 1-4, cells were trypsinized and counted using a Z2 Particle Counter (Beckman Coulter). For all cell proliferation experiments, three wells were used for each experimental condition.

### Oxygen Consumption

Oxygen consumption rates (OCR) and acidification rates (ECAR) of HeLa cells expressing LUC, *Lb*NOX, *Ec*STH, or mito*Ec*STH were measured with the Agilent Seahorse XFe96 Analyzer. Cells were seeded at 4-6 x10^3^ cells per well in Seahorse 96-well cell culture microplates in 80 μL of DMEM^+FBS^. The medium was replaced the next day with 200 μL of DMEM^+dFBS^ ± 300 ng/mL of Dox. Twenty-four hours later, medium was replaced with 200 μL of the Seahorse assay medium [pyruvate free DMEM (US Biological, D9802), 10% dialyzed FBS (LifeTechnologies, 26400-044), and 25 mM HEPES-KOH, pH 7.4]. Basal measurements were collected 6 times and 5 measurements were collected after injection of antimycin A (final concentration 1 μM). For the stress test assay after basal measurements were collected 6 times, oligomycin A, FCCP, and piericidin A + antimycin A were used (injected in this order at a final concentration of 1 μM). Three measurements were taken for oligomycin A and piericidin A + antimycin A and 4 measurements were taken for FCCP. After each assay, the 96-well plate was extensively washed with PBS, and the BCA assay (ThermoFisher, 23227) was used to quantify protein content in each well. This protein concentration was used to normalize data in all Seahorse experiments.

### Determination of Total Cellular NADH/NAD^+^ and NADPH/NADP^+^ Ratios

Two-four hundred thousand cells were seeded in 6-cm dishes in 2 mL of DMEM^+FBS^. Twenty-four hours later, media was exchanged with 3 mL of DMEM^+dFBS^ and 300 ng/mL of Dox. After an additional twenty-four hours, media was exchanged with DMEM^+dFBS^ and 300 ng/mL of Dox that was incubated overnight at 5% CO_2_ or 5% CO_2_/3.5% O_2_ (for IMR-90 human fibroblasts). Three hours later, 6-cm dishes were placed on ice, washed with 3 mL of ice-cold PBS, and lysed with 0.5 mL of 1:1 mixture of PBS and 1% dodecyltrimethylammonium bromide (DTAB) in 0.2 M NaOH. Samples were processed as previously described, transferred to all-white 96-well plates, and luminescence was measured over 1.5 hours using EnVision 2103 or Tecan Infinite F200 PRO plate readers (5). For each experimental condition at least three 6-cm dishes were used, and all assays were performed at least three times.

### Determination of Total Cellular GSH/GSSG Ratios

Five thousand cells were seeded in 0.2 mL of DMEM^+FBS^ in a black 96-well plate with a clear bottom (Corning, 3904). Twenty-four hours later media was exchanged with 0.2 mL of DMEM^+dFBS^ ± 300 ng/mL Dox. After an additional twenty-four hours media was exchanged with DMEM^+dFBS^ ± 300 ng/mL Dox that was incubated overnight at 5% CO_2_. To selected wells 200 μM menadione (vitamin K3) was also added. Three hours later, cells were lysed with 60 μL of either Total Glutathione Lysis Reagent or Oxidized Glutathione Lysis Reagent (provided in Promega GSH/GSSG-Glo Assay kit, V6611). After lysis, 50 μL of sample was transferred to an all-white 96-well plate and topped with 50 μL of Luciferin Generation Reagent. After 30 min incubation, 50 μL of Luciferin Detection Reagent was added to each well. Luminescence was read over 3 hours using Tecan Infinite F200 PRO plate reader. For each cell line or experimental condition, n=3 biological replicates were used.

### Measurements of Reactive Oxygen Species (ROS) in Cells Expressing Luciferase, *LbNOX, Ec*STH and mito*Ec*STH

Six thousand cells were seeded in 0.2 mL of DMEM^+FBS^ in a black 96-well plate with clear bottom (Corning, 3904). Twenty-four hours later media was exchanged with 0.2 mL of DMEM^+dFBS^ ± 300 ng/ml Dox. After an additional twenty-four hours cells were washed in the assay medium [DMEM free from phenol red, pyruvate, folic acid, niacinamide, pyridoxal, riboflavin and thiamine (US Biological, D9800-17), 25 mM HEPES-KOH, pH 7.4, 10% dialyzed FBS (ThermoFisher, 26400044) and 25 mM glucose (ThermoFisher, A2494001)]. CellROX Green Reagent (ThermoFisher, C10444) was added to the assay medium (5 μM final concentration), and cells were then returned to the incubator. After 30 minutes, cells were washed three times in the assay medium, and 200 μM menadione (vitamin K3) or 2 μM antimycin A were added to selected wells. Fluorescence (Ex/Em 485 nm/535 nm) was recorded over 3 hours using a Tecan Infinite F200 PRO plate reader, and only end-point values were used in the data analysis. For each cell line or experimental condition, n=3 biological replicates were used. Additional measurements were taken using MitoSOX Red mitochondrial superoxide indicator (ThermoFisher, M36008). Cells were prepared as stated above with MitoSOX Red added to the assay medium (5 μM final concentration) and incubated for 20 minutes. After incubation, the 96-well plate was washed 3 times with assay medium and 200 μM menadione (vitamin K3) or 2 μM antimycin A were added to selected wells. Fluorescence (Ex/Em 485 nm/580 nm) was recorded over 3 hours using a Tecan Infinite F200 PRO plate reader, and only end-point values were used in the data analysis. For each cell line or experimental condition, n=3 biological replicates were used. For each experiment, an identical control 96-well plate was prepared and used to determine protein content in each well by BCA assay (ThermoFisher, 23227).

### Measurements of H_2_O_2_ Production in Cells Expressing Luciferase, *Lb*NOX, *Ec*STH and mito*Ec*STH

Six thousand cells were seeded in 0.2 mL of DMEM^+FBS^ in a black 96-well plate with a clear bottom (Corning, 3904). Twenty-four hours later, media was exchanged with 0.2 mL of DMEM^+dFBS^ and doxycycline (300 ng/mL). Twenty-four hours after addition of doxycycline, cells were washed in the assay medium [DMEM free from phenol red, pyruvate, folic acid, niacinamide, pyridoxal, riboflavin and thiamine (US Biological, D9800-17), 25 mM HEPES-KOH, pH 7.4, 10% dialyzed FBS (ThermoFisher, 26400044), and 25 mM glucose (ThermoFisher, A2494001)]. To each well containing cells in 50 μL of the assay medium [± 2 μM antimycin, ± 200 μM menadione (vitamin K3)], another 50 μl of solution was added [48.5 μl of the Amplex Red Hydrogen Peroxide/Peroxidase Assay Kit 1X Reaction Buffer (ThermoFisher, A22188), 0.5 μL of Amplex Red (10 mM stock) and 1 μL of HRP (10 U/mL stock)]. Fluorescence (Ex/Em 540 nm/590 nm) was recorded over 3 hours using a Tecan Infinite F200 PRO plate reader and end-point values were used in the data analysis. For each cell line or experimental condition (n=3) biological replicates were used. For each experiment an identical control 96-well plate was prepared and was used to determine protein content in each well using BCA assay (ThermoFisher, 23227).

### Measurements of Lactate and Pyruvate in Spent Media using LC-MS

In the same experiments where cellular NADH/NAD^+^ and NADPH/NADP^+^ ratios were determined, spent media was collected (3 hours after fresh DMEM^+dFBS^ change, see above). Thirty μL of spent media was mixed with 70 μL of icecold acetonitrile and 20 μL of internal standard mix containing: 50 μM ^13^C_3_ pyruvate and 5 mM ^13^C_3_ lactate. Tubes were vortexed for 1 min and then set at −20°C for 20 min. Samples were then centrifuged at 12,700 rpm for 20 min at 4°C, and 100 μL was transferred into autosampler vials for the LC-MS/MS analysis. The samples were run on an Agilent 6495 QqQ with jet stream coupled to an Agilent 1290 LC stack with an Agilent HILIC-Z (2.1×150mm) column (Center for Mass Spectrometry and Metabolomics, TSRI). The mobile phase was composed of A= 10 mM NH_4_OAc, 20 μM deactivator, pH=9 and B= 85:15 ACN/H_2_O 10 mM NH_4_OAc, 20 μM deactivator, pH=9. The gradient started at 100 % B (0-1 min), decreasing to 60 % B (1-5 min) and was followed by an isocratic step (6 min – 10 min) before a 5 min post-run for column re-equilibration. The flowrate was set to 250 μL/min and the sample injection volume was 10 μL. Operating in negative mode, the source conditions were as follows: drying gas temperature set to 200 °C with a flowrate of 11 L/min, sheath gas temp set to 300 °C with a sheath gas flowrate of 12 L/min, neb pressure set to 35 psi, cap voltage set to 3000V, and nozzle voltage set to 0V. Data was processed using Agilent MassHunter Quantitative analysis software. For absolute quantification, calibration curves were generated by comparing the ratio of response of the analyte and its internal standard to its concentration. The linearity of each standard curve was evaluated. Concentrations of compounds were corrected for the ratio of MS response (peak area) between analyte and internal standard to account for matrix affects and any losses during prep.

### Metabolomic Profiling using LC-MS

One-three hundred thousand HeLa Tet-On 3G cells expressing Luciferase, *Lb*NOX, *Ec*STH, and mito*Ec*STH were seeded in a 6-well plate in 2 mL of pyruvate-free DMEM^+FBS^. Twenty-four hours after seeding, media was exchanged to 2 mL of DMEM^+dFBS^ ± 300 ng/ml Dox. After an additional twenty-four hours, cells were removed from the incubator, placed on ice, lysed (without a PBS wash) with 1 mL of dry-ice cold 80% methanol/20% water solution containing 1.5 μM Cambridge AA mix (Cambridge Isotope Laboratories, MSK-A2-1.2). Immediately after, 6-well plates were transferred to −80°C and incubated overnight to help protein precipitation and allow equilibrium between solubilized metabolites and precipitated protein. The 6-well plates were scraped the next day, and all material was transferred to 1.5 mL Eppendorf tubes and dried in a SpeedVac.

For metabolomic profiling using liquid chromatography–mass spectrometry, dried extracts were resuspended in 60 μL of water for ion pair liquid chromatography separation. Samples were vortexed, incubated on ice for 20 min and clarified by centrifugation at 20,000*g* for 20 min at 4 °C. Ion pair LC–MS analysis was performed on a 6230 TOF mass spectrometer (Agilent Technologies) in negative ionization mode using a XSelect HSS T3, 150 x 2.1 mm, 3.5 μm particle size (Waters), and using a gradient of solvent A (5 mM octylamine and 5 mM acetic acid in water) and solvent B (5 mM octylamine and 5 mM acetic acid in methanol), with a post-column flow consisting of a blend of acetone:DMSO (90:10) at 300 μL/min. The analytical gradient was 0-3.5 min, 1% B; 4-15 min, 35% B; 20-22 min, 100% B; 22-27 min, 1% B. Other LC parameters were as follows: flow rate 300 μL/min, column temperature 40°C, and injection volume 5 μL. MS parameters were as follows: gas temp: 250°C; gas flow: 9 L/min; nebulizer pressure: 35 psi; sheath gas temp: 250°C; sheath gas flow: 12 L/min; VCap: 3500 V; and fragmentor: 125 V. Data were acquired from 50 to 1700 m/z with active reference mass correction (m/z: 119.0363 and 966.0007) infused through a second nebulizer according to the manufacturer’s instructions. Peak identification and integration were done based on exact mass and retention time match to commercial standards. Data analysis was performed with MassHunter Profinder software v10.0 (Agilent Technologies) and Skyline v22.2.

### Expression and Purification of Recombinant *Ec*STH

*E. coli* BL21 (DE3) cells (Life Technologies, C6010-03) were transformed with the pET30a-Hisx6-*Ec*STH-FLAG vector and were grown at 37°C in six 2.8-L flasks, each containing 1 L of Terrific Broth (TB) medium supplemented with 50 μg/mL kanamycin. When absorbance at 600 nm reached 0.4-0.6, the temperature was decreased to 15°C, and cells were grown for additional 2 hours before protein expression was induced with 0.1 mM isopropyl β-D-1-thiogalactopyranoside (IPTG). Recombinant *E. coli* STH was purified using a protocol previously described for *L. brevis* H_2_O-forming oxidase (*Lb*NOX) with several modifications (see below) (5). More specifically, the cell pellet was resuspended in 350 mL of the lysis buffer (50 mM Na_2_HPO_4_, pH 8.0, 500 mM NaCl and 30 mM imidazole), and affinity chromatography was performed using an Omnifit glass column packed with 35 mL of Ni Sepharose 6 Fast Flow (Cytiva). For the Source 15Q anion-exchanger step, protein was eluted with the gradient of 50-350 mM NaCl in 50 mM Na_2_HPO_4_, pH 7.5. The most active STH fractions were pooled and subjected to size-exclusion chromatography on a HiLoad 16/600 Superdex 200 (Cytiva, 28989335) equilibrated with 50 mM HEPES-NaOH pH 7.5 and 150 mM NaCl (buffer A). Resulting fractions were pooled, concentrated, flash-frozen in liquid nitrogen, and stored at −80 °C.

### Determination of the Oligomerization State of Recombinant *Ec*STH

Analytical size-exclusion chromatography was performed using a Superdex 200 Increase 10/300 GL column (Cytiva, 28990944) equilibrated in buffer A. A calibration curve was produced using thyroglobulin (669 kDa), ferritin (440 kDa), beta amylase from sweet potato (200 kDa), aldolase (158 kDa), conalbumin (75 kDa), bovine albumin (66 kDa) and ovalbumin (44 kDa). Protein samples were injected with a gastight micro-syringe and the column was operated at a flow rate of 0.9 mL/min.

### UV-Visible Spectroscopy and Activity Assays of Recombinant *Ec*STH

UV-visible spectra were recorded on a Cary 3500 spectrophotometer (Agilent, CA). *Ec*STH (10–20 μM FAD active sites) in buffer A was incubated at 24 °C, and 3 mM sodium dithionite was added under aerobic conditions. An extinction coefficient (ε450 =11.3 mM^-1^cm^-1^) was used to quantify FAD. Activity of *Ec*STH was monitored in the presence of thio-NAD^+^ (Sigma, T7375) or thio-NADH (VWR, TCT2980) by following of absorbance at 398 nm (ε_398_ =11.3 mM^-1^cm^-1^). Michaelis–Menten analysis was performed by varying one substrate (5–1000 μM) while a second co-substrate was kept constant at 200 μM as indicated in (**Fig. 2B-C** and **Fig.S2A-D**). Substrates were fixed at 200 μM due to substrate inhibition (14,55). A reaction mixture containing 0.2 mL of buffer A was incubated for 3 min at 37 °C before both substrates and enzyme (0.3-0.6 μg) were added. The multiwavelength option of Cary 3500 spectrophotometer (Agilent, CA) allowed to simultaneous monitoring both 340 and 398 wavelengths (only absorbance at 398 nm was used for quantification). The *k*_cat_ values for *Ec*STH were calculated per monomer of the protein.

### RNA-Seq of HeLa cells Expressing Luciferase, *Lb*NOX, *Ec*STH and mito*Ec*STH

Five hundred thousand cells were seeded in 10 cm dishes in 10 mL of DMEM^+FBS^. Twenty-four hours later, media was exchanged with 10 mL of DMEM^+dFBS^ ± 300 ng/mL Dox. Twenty-four hours after addition of doxycycline, cells were harvested, and cell pellets were snapfrozen. Samples were submitted to GENEWIZ for NGS RNA sequencing. The integrity of RNA was analyzed on Agilent Bioanalyzer, based on the integrity of the 18S and 28S ribosomal sequences of mammalian genomes. All groups passed with an RNA integrity number (RIN) value of 9 or above. All RNA Libraries from each group were sequenced on an Illumina HiSeq X instrument, generating 2x 150 bp paired end reads. Using Trimmomatic v.0.36 sequence reads were trimmed to remove possible adapter sequences and nucleotides with poor quality. Using the STAR aligner v.2.5.2b., the trimmed reads were mapped to the *Homo sapiens* GRCh38 reference genome available on ENSEMBL. The STAR aligner is a splice aligner that detects splice junctions and incorporates them to align the entire read sequences. Unique gene hit counts were calculated by using feature Counts from the Subread package v.1.5.2. The hit counts were summarized and reported using the gene_id feature in the annotation file. Only unique reads that fell within exon regions were counted. If a strand-specific library preparation was performed, the reads were strand-specifically counted. Counts were then processed in R using the DESeq2 package (61). Gene log2 fold changes were calculated based on log2(Group 2 mean normalized counts/Group 1 mean normalized counts) as implemented in DESeq2. P-values were derived from the Wald test and adjusted using the Benjamini and Hochberg method. Genes with an adjusted p-value < 0.05 and absolute log2 fold change >1 were recognized as differentially expressed genes. Using ggplot2 packages in R, volcano plots were generated to display the global differentially expressed genes across the compared groups. Each data point in the scatter plot represents a gene. The log2 fold change of each gene is represented on the x-axis and the minus log10 of its adjusted p-value is represented on the y-axis. Genes with an adjusted p-value less than 0.05 and a log2 fold change greater than 1 are indicated by red dots, representing up-regulated genes. Genes with an adjusted p-value less than 0.05 and a log2 fold change less than −1 are indicated by blue dots, representing down-regulated genes. Genes without significant changes are indicated by gray dots. The most significantly changed genes are named in each volcano plot. Gene ontology clusters were formed using comparisons of significantly changed genes in *Ec*STH expressing cells to those in luciferase expressing cells, and the enrichment of gene ontology terms was tested using Fisher exact test (GeneSCF v1.1-p2). Significantly changed gene sets were defined by a Benjamini-Hochberg adjusted p-value <0.05, differentially expressed gene sets (up to 20 terms) were shown in a bar graph. The individual genes from the top 2 differentially expressed gene sets, the GO:0055114~oxidation-reduction process and GO:0006695~cholesterol biosynthetic process, were highlighted in the volcano plot in different colors. The overlap of these two gene sets is presented in a Venn diagram.

### Statistical Analysis and Reproducibility

For repeated measurements, a Welch ANOVA test with unpaired t test was performed using a built-in statistical package in GraphPad Prism 9.3.1. All experiments were performed at least three times with similar results. P values are denoted as: not significant [ns], p<0.05 [*], p<0.01 [**], p<0.001 [***], p<0.0001 [****]. Each dot in a bar graph represents a biological replicate. All error bars displayed in figures show standard deviation (S.D.).

## ACKNOWLEDGMENTS

We thank members of the Thompson laboratory at MSKCC, Michelle Clasquin, Amy Caudy and Adam Rosebrock for discussion and feedback. We thank the Scripps Center for Metabolomics for their assistance with metabolomics services. This work was supported by grants from the National Institutes of Health (R00GM121856, R03AG067301 and R35GM142495 to VC). The Seahorse XFe96 analyzer in the Saez laboratory (TSRI) was supported by 1S10OD16357.

## CONTRIBUTIONS

VC conceived the study. MLH, ENA and ALZ performed all experiments with assistance from XP and CY. CY, MLH and ENA performed kinetic characterization experiments. SV and JRC performed LC-MS experiments. XP analyzed RNA-Seq data. VC and MLH wrote the manuscript with input from all the authors. All authors contributed to editing the manuscript and approved the final version.

---

**Supplementary Figure S1.**
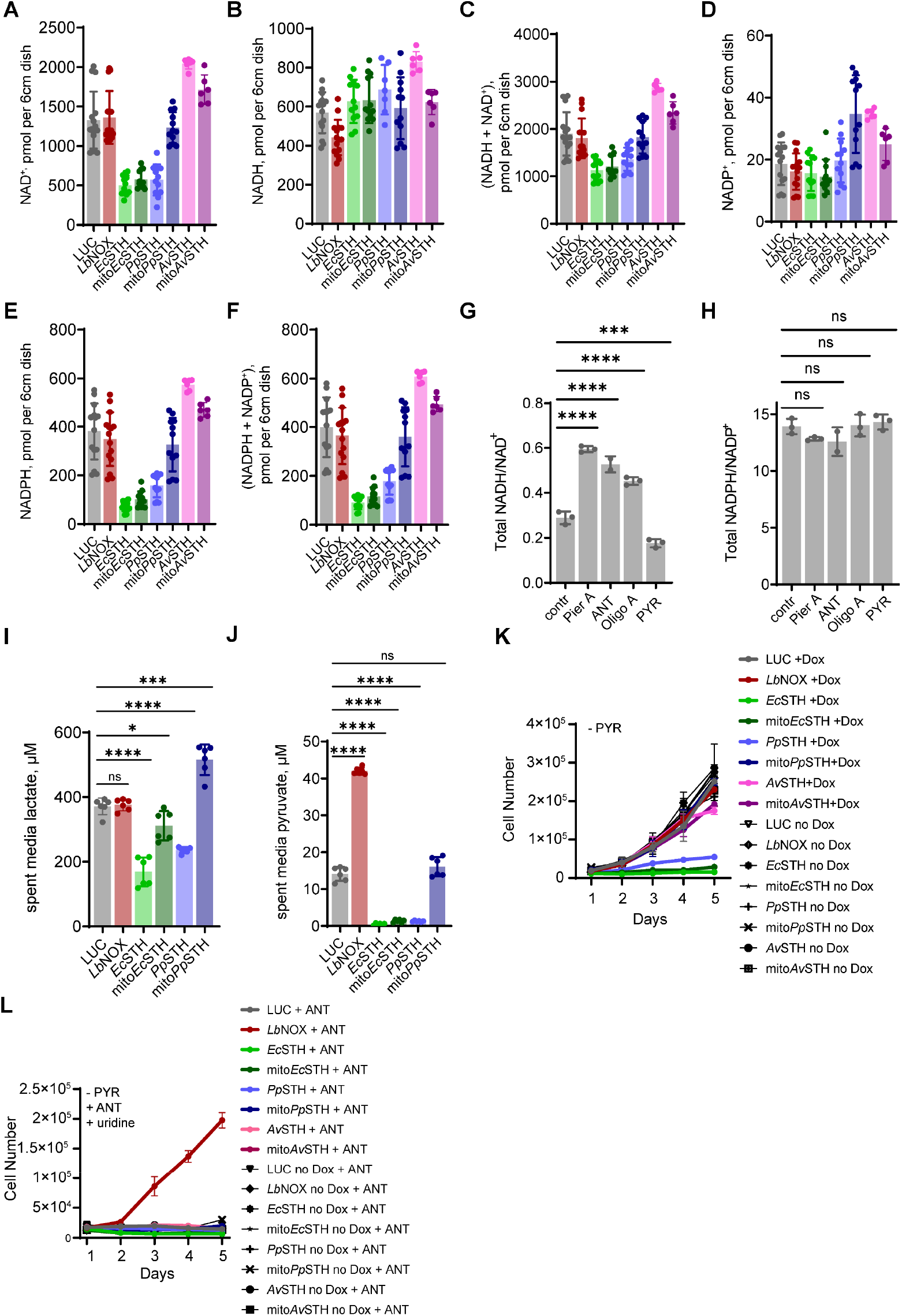
Screening of bacterial STHs in HeLa cells. Effect of STHs expression from *E. coli, A. vinelandii*, and *P. putida* on amounts of NAD^+^ (**A**), NADH (**B**), total NAD pool (NAD^+^ + NADH) (**C**), NADP^+^ (**D**), NADPH (**E**) and total NADP pool (NADP^+^ + NADPH) (**F**). The values correspond to ratios in Fig. 1C-D. The total NADH/NAD^+^ **(G)** and NADPH/NADP^+^ **(H)** ratios measured in *wild-type* HeLa cells three hours after changing to fresh pyruvate-free DMEM^+dFBS^ with 1 μM antimycin A (ANT), 1 μM piericidin A (Pier), 1 μM oligomycin A (Oligo A), or 1 mM pyruvate (PYR). Lactate (**I**) and pyruvate (**J**) levels in media which was incubated for 3 hours (spent media) with HeLa cells expressing untargeted and mitochondrially targeted STHs from *E. coli* and *P. putida*. The effect of expression of untargeted and mitochondrially targeted STHs from *E. coli, P. putida* and *A. vinelandii* on proliferation of HeLa cells in pyruvate-free DMEM^+dFBS^ with and without 300 ng/mL Dox (**K**). In (**L**) pyruvate-free DMEM^+dFBS^ was supplemented with 1 μM antimycin A (ANT) and 200 μM uridine. LUC and *Lb*NOX expressing HeLa cells were used as controls in (A-F) and (I-L). Statistical significance indicated for (G-J) represents a Welch ANOVA test with an unpaired t test. For growth curves (K-L), error bars represent S.D. based on n=3 replicates per time point per experimental condition. All growth curve experiments were repeated at least (n=3) times. p<0.05*, p<0.01**, p<0.001**, p<0.0001****

**Supplementary Figure S2.**
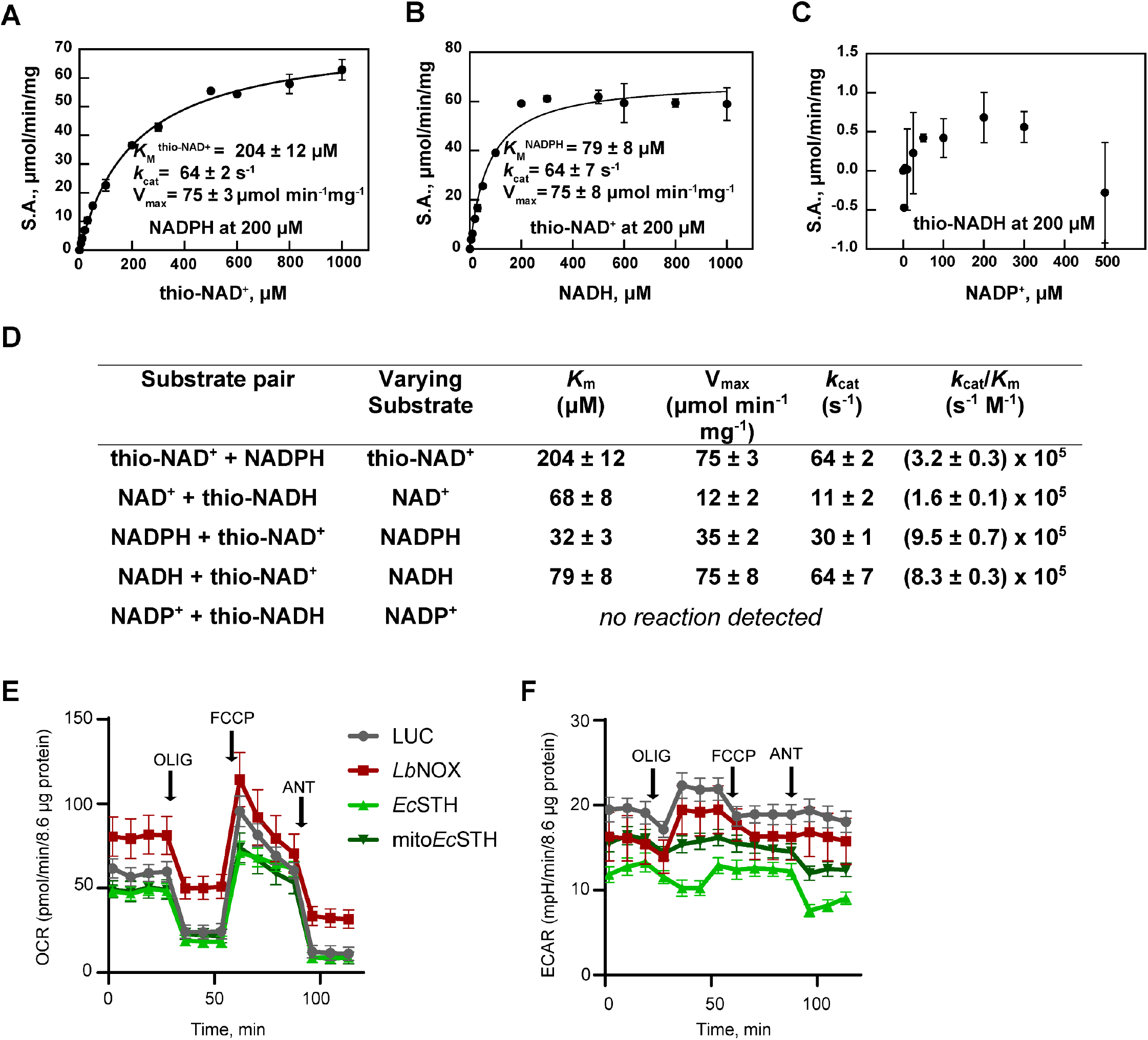
Biochemical properties of recombinant *E. coli* STH (*Ec*STH) and bioenergetics of *Ec*STH and mito*Ec*STH expression in HeLa cells. **(A-B)** Michaelis-Menten analysis of the reaction catalyzed by *Ec*STH with thio-NAD^+^ (A) or NADH (B). **(C)** No reaction was observed for NADP^+^ and thio-NADH substrate pair. Reported values for V_max_, *k*_cat_ and *K*_M_ represent the mean ± S.D. from (n=3) independent experiments. **(D)** Summary of kinetic parameters for all substrate pairs tested in this study (see also Fig. 2B-C). *K*_cat_ values were calculated per monomers of FAD active sites. (**E-F**) mitochondrial stress test where OCR and ECAR were measured before and after sequential injection of 1 μM oligomycin A (OLIG), 2 μM FCCP, and 2 μM antimycin A (ANT). LUC and *LbNOX* expressing HeLa cells were used as controls in (E-F).

**Supplementary Figure S3.**
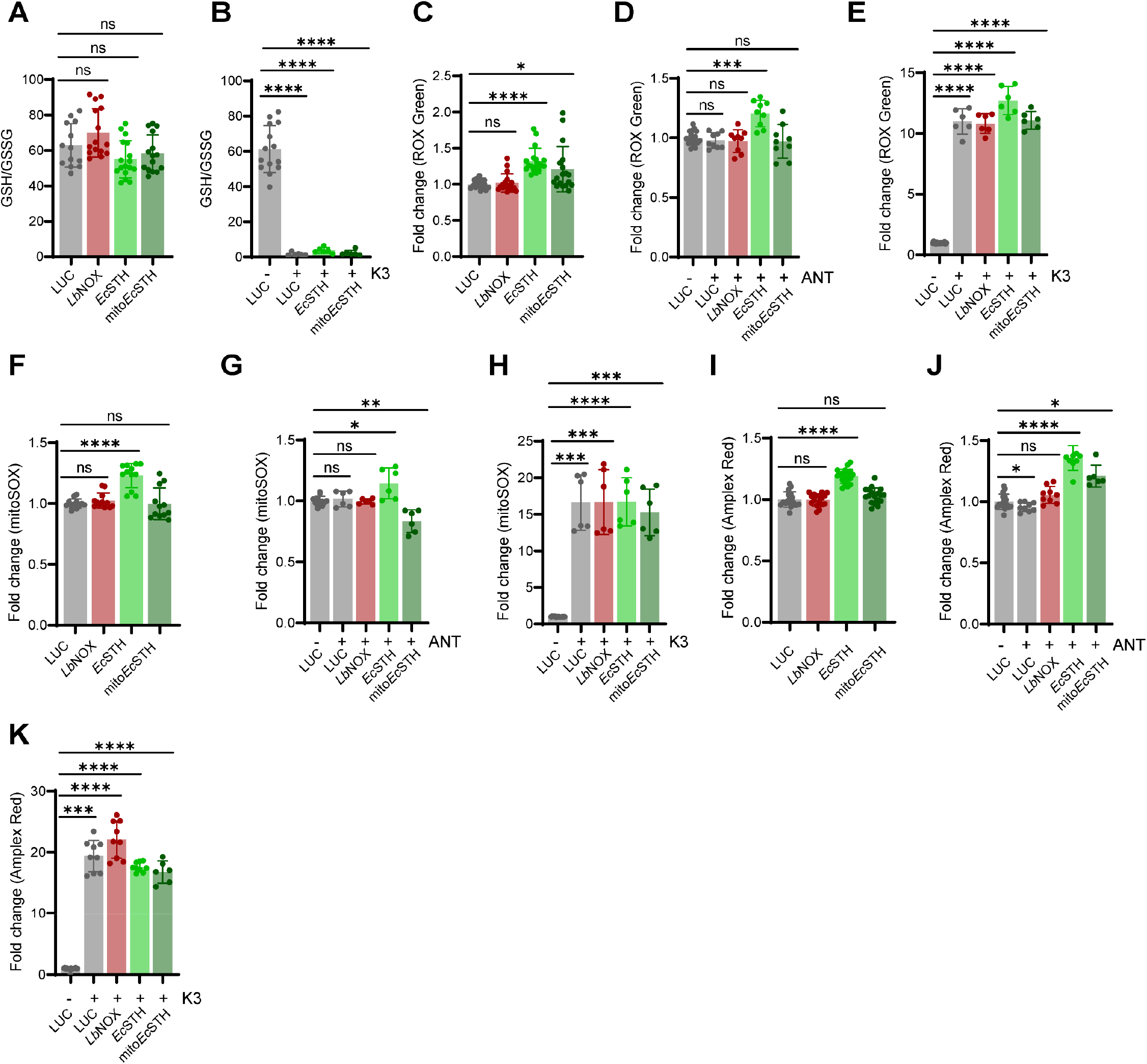
Effect of *Ec*STH and mito*Ec*STH expression in HeLa cells on GSH/GSSG ratios and reactive oxygen species (ROS) levels. The GSH/GSSG ratios **(A-B)**, ROS levels determined with CellROX Green **(C-E)**, mitoSOX **(F-H)**, and H_2_O_2_ levels determined with Amplex Red **(I-K)** in HeLa cells expressing *Ec*STH and mito*Ec*STH. Two hundred μM menadione (K3) or 1 μM antimycin A (ANT) were used as positive controls in (B, D, E, G, H, J and K). LUC and *Lb*NOX expressing HeLa cells were used as controls in (A-K). Statistical significance indicated for (A-K) represents a Welch ANOVA test with an unpaired t test. p<0.05*, p<0.01**, p<0.001**, p<0.0001****

**Supplementary Figure S4.**
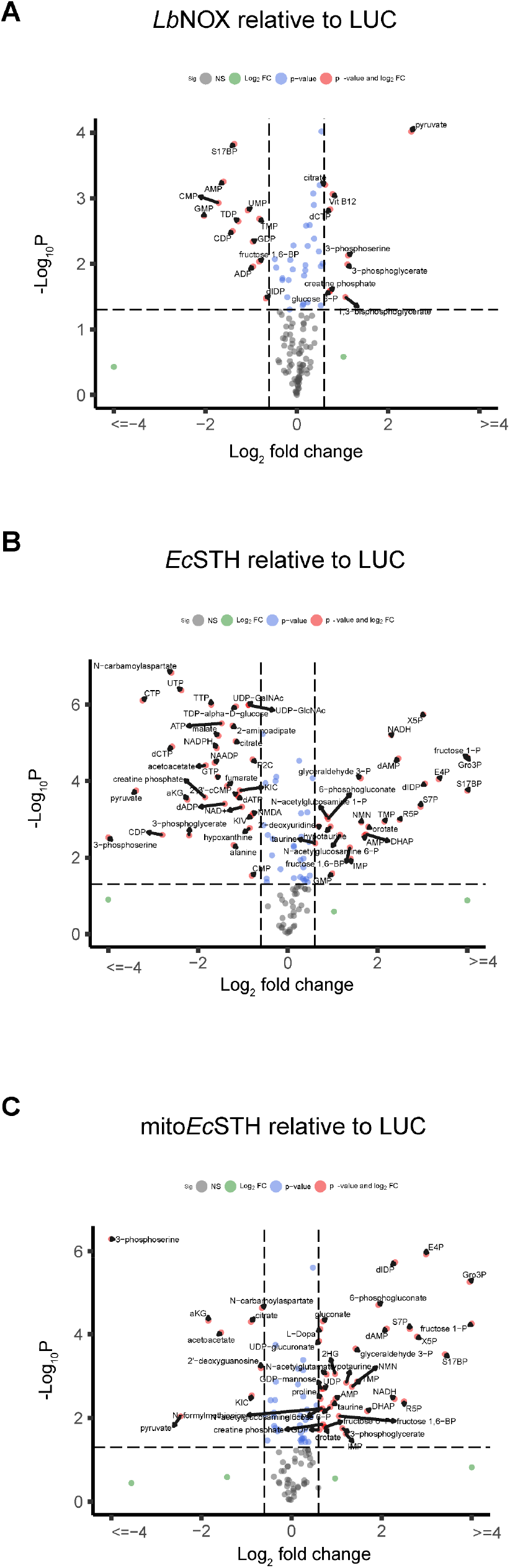
Metabolic features of *LbNOX, Ec*STH and mito*Ec*STH expression in HeLa cells. Targeted metabolomics in HeLa cells expressing *Lb*NOX (**A**), *Ec*STH (**B**) and mito*Ec*STH (**C**). LUC expressing HeLa cells were used as a background in (A-C). The statistical significance indicated for (A-C) represents p value cutoff = 0.05, fold change cutoff = 1.5 and Welch t test (unadjusted).

**Supplementary Figure S5.**
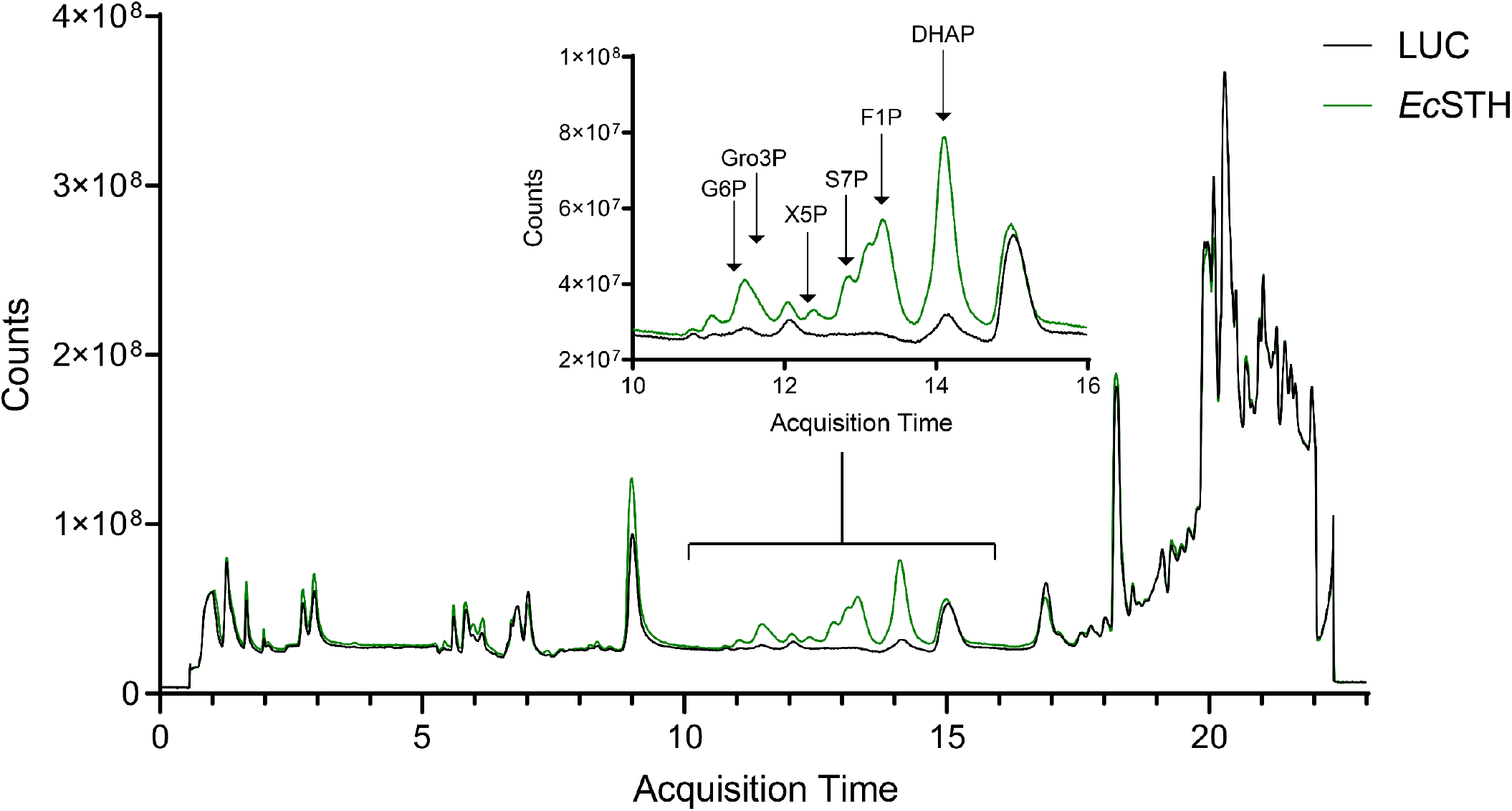
Reductive stress leads to accumulation of non-canonical sugar phosphates. Total ion chromatograms (TICs) of representative LUC and *Ec*STH samples. *Ec*STH expression leads to increased signal intensity specifically in the chromatographic region where sugar phosphates and other glycolysis intermediates elute. Peaks which we were able to annotate are labeled. G6P: glucose-6-phosphate; Gro3P: glycerol 3-phosphate; X5P: xylulose-5-phosphate; S7P: sedoheptulose-7-phosphate; F1P: fructose-1-phosphate; DHAP: dihydroxyacetone phosphate.

**Supplementary Figure S6.**
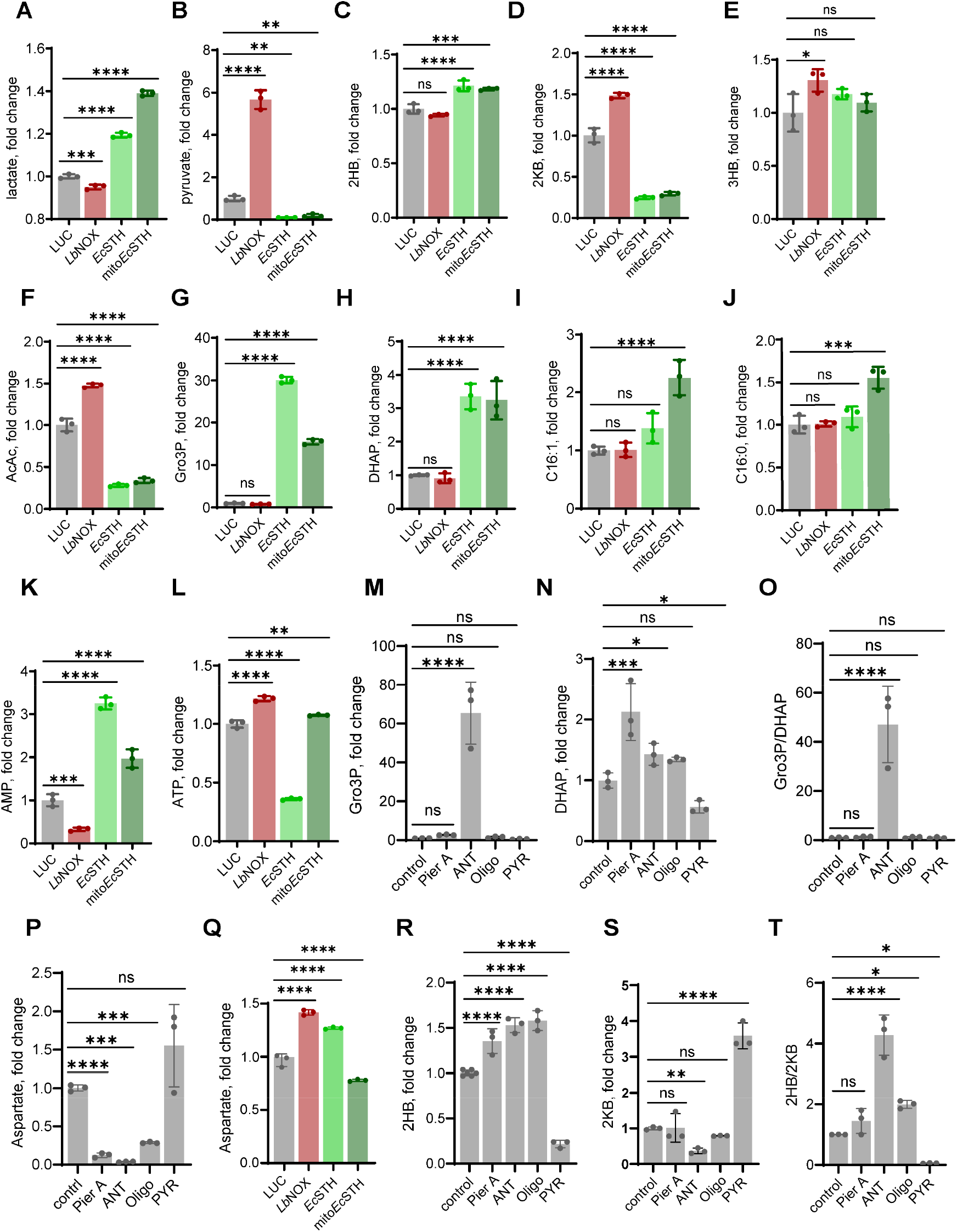
Levels of various metabolites in HeLa cells expressing *Ec*STH and mito*Ec*STH. Lactate (**A**), pyruvate (**B**), 2HB (**C**), 2KB (**D**), 3HB (**E**), AcAc (**F**), Gro3P (**G**), DHAP (**H**), C16:1 (**I**), C16:0 (**J**), AMP (**K**) and ATP (**L**) levels in HeLa cells expressing *Ec*STH and mito*Ec*STH. Gro3P (**M**), DHAP (**N**), Gro3P/DHAP ratio (**O**) and aspartate (**P**) levels in *wild-type* HeLa cells three hours after changing to fresh pyruvate-free DMEM^+dFBS^ with 1 μM piericidin A (Pier A), 1 μM antimycin A (ANT), 1 μM oligomycin A (Oligo), or with 1 mM pyruvate (PYR). Aspartate levels in HeLa cells expressing *Ec*STH and mito*Ec*STH (**Q**). 2HB (**R**), 2KB (**S**) and 2HB/2KB ratio (**T**) in *wild-type* HeLa cells three hours after changing to fresh pyruvate-free DMEM^+dFBS^ with 1 μM piericidin A (Pier A), 1 μM antimycin A (ANT), 1 μM oligomycin A (Oligo A), or with 1 mM pyruvate (PYR). LUC and *Lb*NOX expressing HeLa cells were used as controls in (A-L and Q). Statistical significance indicated for (A-T) represents a Welch ANOVA test with an unpaired t test. p<0.05*, p<0.01**, p<0.001**, p<0.0001****.

**Supplementary Figure S7.**
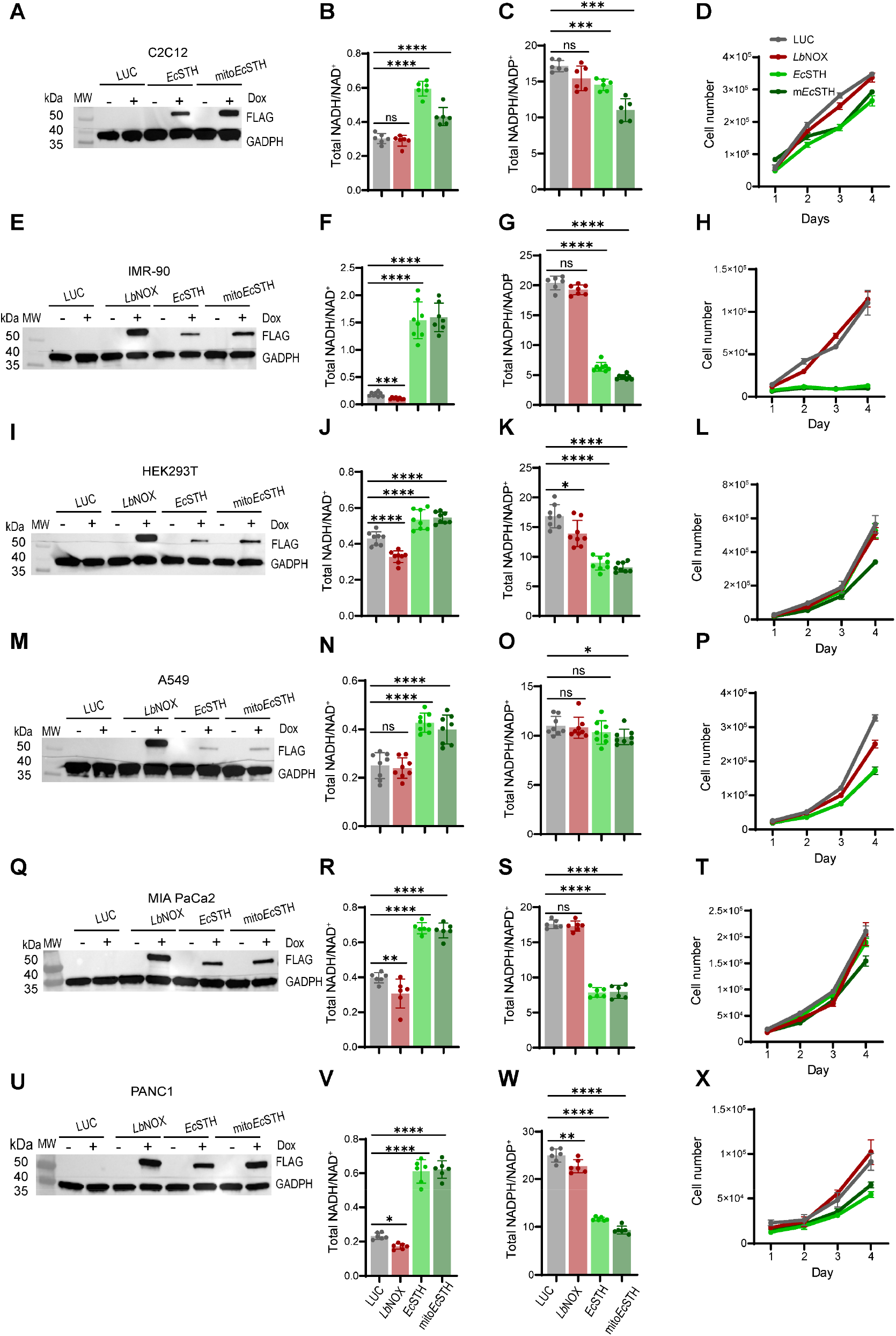
The anti-proliferative effect of *Ec*STH and mito*Ec*STH expression is cellular background specific. Western blot analysis of *Ec*STH and mito*Ec*STH expression in C2C12 mouse myoblasts **(A)**, IMR-90 human fibroblasts **(E)**, embryonic human kidney HEK293T cells **(I)**, A549 lung adenocarcinoma cells **(M),** MIA PaCa-2 epithelial tumor tissue of the pancreas **(Q)** and PANC-1 pancreatic duct epithelioid carcinoma **(U)**. The total NADH/NAD^+^ ratios in C2C12 myoblasts **(B)**, IMR-90 **(F)**, HEK293T **(J)**, A549 **(N)**, MIA PaCa-2 **(R)** and PANC-1 **(V)** cells expressing *Ec*STH and mito*Ec*STH. The total NADPH/NADP^+^ ratios in C2C12 myoblasts **(C)**, IMR-90 **(G)**, HEK293T **(K)**, A549 **(O)**, MIA PaCa-2 **(S)** and PANC-1 **(W)** cells expressing *Ec*STH and mito*Ec*STH. Proliferation of C2C12 myoblasts **(D)**, IMR-90 **(H)**, HEK293T **(L)**, A549 **(P)**, MIA PaCa-2 **(T)** and PANC-1 **(X)** cells expressing *Ec*STH and mito*Ec*STH in pyruvate-free DMEM^+dFBS^. LUC and *Lb*NOX expressing HeLa cells were used as controls in (A-X). The statistical significance indicated for (B-C, F-G, J-K, N-O, R-S, V-W) represents a Welch ANOVA test with an unpaired t test. p<0.05*, p<0.01**, p<0.001**, p<0.0001****. For growth curves (D, H, L, P, T and X) error bars represent S.D. based on n=3 replicates per time point per condition. All cellular proliferation experiments were repeated at least n=3 times.

